# Renalase inhibition regulates β cell metabolism to defend against acute and chronic stress

**DOI:** 10.1101/2024.06.11.598322

**Authors:** Tara MacDonald, Birgitta Ryback, Jessica Aparecida da Silva Pereira, Siying Wei, Bryhan Mendez, Erica Cai, Yuki Ishikawa, Gordon Weir, Susan Bonner-Weir, Stephan Kissler, Peng Yi

## Abstract

Renalase (Rnls), annotated as an oxidase enzyme, is a GWAS gene associated with Type 1 Diabetes (T1D) risk. We previously discovered that Rnls inhibition delays diabetes onset in mouse models of T1D *in vivo*, and protects pancreatic β cells against autoimmune killing, ER and oxidative stress *in vitro*. The molecular biochemistry and functions of Rnls are entirely uncharted. Here we find that Rnls inhibition defends against loss of β cell mass and islet dysfunction in chronically stressed Akita mice *in vivo*. We used RNA sequencing, untargeted and targeted metabolomics and metabolic function experiments in mouse and human β cells and discovered a robust and conserved metabolic shift towards glycolysis, amino acid abundance and GSH synthesis to counter protein misfolding stress, *in vitro*. Our work illustrates a function for Rnls in mammalian cells, and suggests an axis by which manipulating intrinsic properties of β cells can rewire metabolism to protect against diabetogenic stress.

## Introduction

Type 1 (T1D) and Type 2 Diabetes (T2D) are both characterized by inadequate β cell mass, β cell dysfunction, and the eventual development of hyperglycemia across a heterogeneous timescale as the β cell fails^1^. The loss of glucose-stimulated insulin secretion predicts both T1D and T2D as an early signal, but the routes to β cell demise differ ^2–5^. In T1D, β cell death stems from innate and adaptive immune-mediated destruction of pancreatic β cells, directly by perforin/granzyme activities^6^, and less directly by cytokine-driven injury^7–12^. In contrast, immune contributions to β cell damage in T2D are less clear. Deposits of amyloid, which are aggregates of the β cell-secreted product islet-associated polypeptide (IAPP), are often found in T2D and damage is thought to be caused by toxic oligomers that develop as amyloid aggregates are formed^13–15^. In T2D, β cell dysfunction develops alongside deteriorations in peripheral insulin sensitivity. Insulin resistance associated with T2D places extra demand on β cells, which can trigger intrinsic endoplasmic reticulum (ER) and oxidative stress^16^. These pathways may act in vicious cycles to synergistically aggravate β cell dysfunction and loss of β cell mass (reviewed in^17^). The key roles for autoimmunity in T1D and insulin resistance in T2D are canonical. In each situation, β cells face the damaging effects of distinct stressors. However, it is now thought that as β cell mass and function fall in T1D, the stress exerted by rising glucose levels may closely resemble the damage that occurs during T2D. In both types of diabetes, β cells may be exposed to similar ER stress, oxidative stress and hyperglycemia, which each impact β cell secretory function and vulnerability.

β cell stress has re-emerged in the spotlight as a pathophysiological driver of both T1D and T2D. The relative contributions of stress mechanisms to pathophysiology are unclear^18–20^. Conceptually, “stress” is considered an aggregate measure of strength and duration of a perturbation that shifts a cell away from homeostasis. Biologically, stress phenomena are less well-defined, especially in the transition from acute signalling‒often natural or even hormetic‒ to chronic (mal)adaptations in the context of disease^17^. Several β cell features impart remarkably high vulnerability to stress relevant to both T1D and T2D: β cell metabolism is regulated by rate-limiting, low Km glucokinase and near-constant glucose flux through mitochondrial metabolic and bioenergetic pathways that can generate reactive oxygen species (ROS)^21^; the ER of β cells works at a high biosynthetic rate to package and secrete proteins, including but not limited to insulin, which can generate substantia ER stress^22,23^ and frequent protein misfolding and activation of the unfolded protein (UPR) and/or integrated stress response (ISR) pathways can disrupt transcription and translation during T1D and T2D^24,25^. In both T1D and T2D, β cell demise is likely driven by a constellation of stressors. It then follows that cell-based therapies – as standalone strategies or adjuncts to immunotherapy – will be essential for protecting β cell mass and function as both deteriorate in natural T1D and T2D, and for shielding stem cell-derived β cells (sc-β cells) in the context of transplantation for replenishing β cell mass^18,26^.

We previously harnessed genome-wide CRISPR screening and selective autoimmune pressure to identify mutations within β cells that might protect against their destruction^27^. Amongst 11 putative protective targets, the Renalase (*Rnls*) gene associated with T1D overall risk and age of onset in GWAS and justified its targeting^28,29^. Using mouse β cell lines and human sc-β cells, we validated that genetic or pharmacological Rnls inhibition conferred protection against T cell killing *in vitro* and *in vivo*, in part by reducing β cell ER and oxidative stress^27^. These findings support the notion that reducing intracellular stress can protect against autoimmune-directed β cell dysfunction and death. Despite these strong phenotypes, the molecular mechanisms and biochemistry of Rnls and its inhibition are entirely unknown.

This study was sparked by the idea that elucidating the function(s) of Rnls would yield key information about broader molecular networks that protect β cells from diabetogenic demise. Here we discovered that Rnls inactivation reprograms fundamental cell metabolism pathways towards steady-state glycolysis and glutathione (GSH) synthesis to protect against hyperglycemia, ER and oxidative stress, on acute to chronic timescales *in vitro* and *in vivo*. This work bolsters the concept of manipulating β cell properties to reduce dysfunction and death during T1D and/or T2D.

## Results

### Pargyline treatment improves glycemia and β cell function in the Akita mouse model

Using structure-based modelling, we previously identified Pargyline (PG) as a repurposed FDA-approved drug that dose-dependently inhibits Rnls from 5µM^27^. We showed a protective effect of PG *in vivo* in models of diabetes – including pro-survival of NIT-1 β cells transplanted into non-obese diabetic (NOD) mice, and reduced diabetes incidence in multiple low dose streptozotocin (STZ)-injected mice – explained at least in part by resistance to acute ER stress and dampened ER stress signal transduction^27^. Here we used the Akita mouse model to test whether *in vivo* stress protection is conferred by PG on a chronic timescale, in absence of autoimmunity as a confounding factor. The Akita strain harbours a point mutation in *Ins2* that results in disrupted disulfide bond formation and misfolded insulin protein, downstream ER stress, loss of β cell mass and hyperglycemia by weaning age (Extended Data 1A)^30–32^. Thus, in absence of autoimmune mechanisms, we conclude that the Akita resembles T1D and T2D in the biology of β cell stress.

We administered 25µg/mL oral PG in drinking water from day 20 (d20) and measured indices of glucose homeostasis in littermate Akita, Akita + PG and WT controls 4 weeks later (Fig 1A). Male mice showed a partial reversal of fasting hyperglycemia (Fig 1B) after 4 weeks, accompanied by a ∼10%, yet insignificant, increase in circulating insulin (Fig 1C). The typical loss of β cell mass in male Akita mice was not spared by PG treatment (Extended Data 1B). Hyperglycemia was quite exacerbated in Akita males compared to less-characterized females (Fig 1B versus Fig 1E). We seized this as an opportunity to use both sexes as models of physiological versus supraphysiological hyperglycemia provoked by the insulin misfolding mutation. Since PG has some degree of off-target effect on monoamine oxidase (MAO) enzymes and particular sensitivity for the B isoform (MAO-B), and because we were unsure of sex differences in PG pharmacodynamics, we collected liver from mice treated with 0-25μg/mL PG in drinking water for 7 days to measure MAO-B activity. Females had considerably higher basal activity, were far less responsive to PG-mediated inhibition and sustained 2x the MAO-B signal compared with males at identical PG doses (Fig 1D). Based on this observation, we doubled the dose of PG and repeated the same study design. Only at 50µg/mL was PG able to reverse hyperglycemia in females to parallel WT controls (Fig 1E), alongside a concomitant increase in circulating insulin (Fig 1F). β cell mass within PG-treated female Akita mice did not differ from the WT controls (Fig 1G, 1H). However, Akita males did not benefit from increasing PG to 50µg/mL (Fig 1B, 1C). Improvements in glycemia were not explained by glucose tolerance or peripheral insulin sensitivity (Extended Data 1C-1F), suggesting improved glycemia as a result of β cell function as opposed to peripheral glucose or insulin handling. Since PG is not 100% specific to Rnls, some of the *in vivo* effects of PG may be driven by off-target drug mechanisms. An early report on MAO inhibitors reported an effect of high dose PG on insulin secretion potentially unrelated to MAO activity^33^. MAO-B is present in islets and in β cells, specifically, but its abundance in the pancreas is less than other tissues (i.e. liver, adipose tissue, skeletal muscle; Human Protein Atlas)^34^.

**Figure 1:**
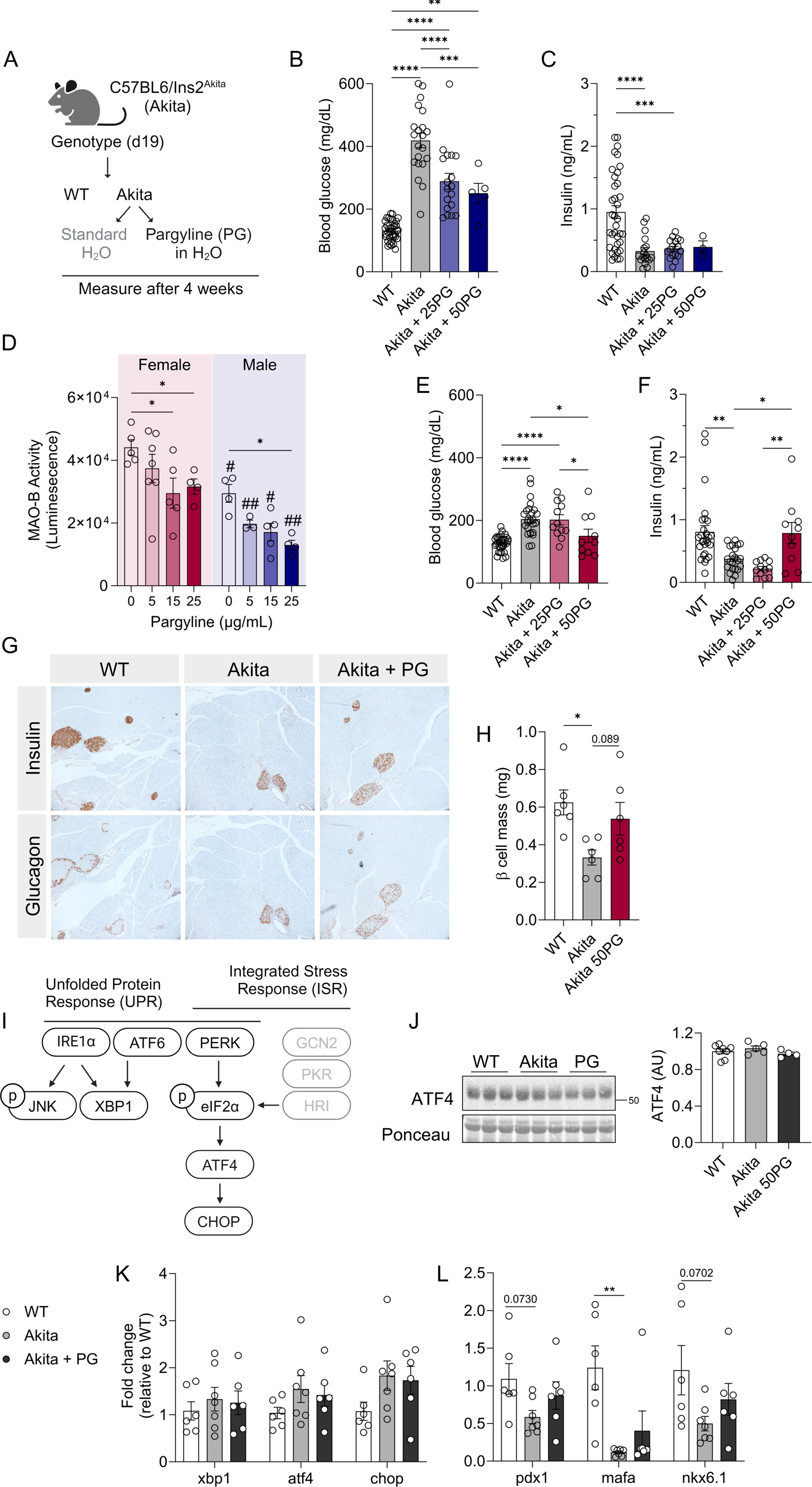
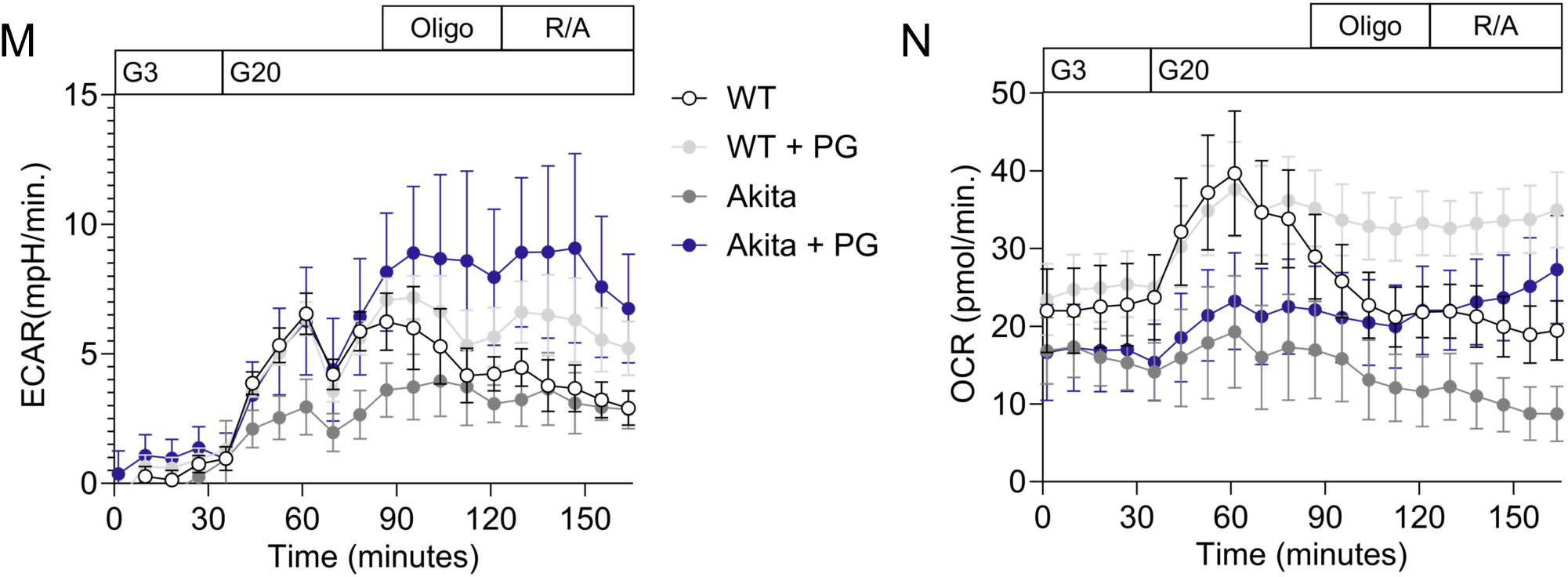
Pargyline improves glycemia in chronically ER-stressed Akita mice. (A) Experiment design: Mice (male and female) were genotyped at day 19 (d19) and allocated to wildtype (WT, untreated), Akita (untreated) or Akita + Pargyline (PG, 25-50µg/mL) at day 20 for 4 weeks. (B) 12h fasting blood glucose and (C) plasma insulin in male mice; n= 3-35/group. (D) Monoamine oxidase B (MAO-B) activity was measured in liver from female (red) and male (blue) mice treated 1week with 0-25 µg/mL PG in drinking water; n=4-7/group. (E) 12h fasting blood glucose and (F) plasma insulin in female mice; n= 10-29/group. (G) Insulin and glucagon staining in female mice with quantification of β cell mass in (H), n= 6/group. (I) Schematic indicating canonical ER stress response pathways and effectors in the UPR and ISR. (J) Islets isolated from female Akita mice were treated with PG (5µM) for 2 days *in vitro* to measure ATF4 protein; n=4-8/group. (K) qPCR measurement of ER stress and (L) maturity markers in islets isolated from Akita mice treated with 50 µg/mL PG for 7 days *in vivo* (n=6-7/group). Islets from a separate cohort of Akita mice treated with 50 µg/mL PG for 7 days were used for metabolic function experiments shown as Extracellular acidification (ECAR; M) and (N) Oxygen consumption (OCR); n= 5-7mice/group. Data is expressed as mean ± SEM. Statistical significance *p ≤ 0.05; ** p ≤ 0.01; *** p ≤ 0.001; *** p ≤ 0.0001 as indicated. Each dot represents an individual mouse. Abbreviations: PG, pargyline; G3, 3mM glucose; G20, 20mM glucose; Oligo, oligomycin; R/A, rotenone/antimycin A.

Presumably the mechanisms of PG in Akita islets *in vivo* would recapitulate ER stress protection observed in β cells *in vitro*. Typical stress pathways, including the ER-stress activated unfolded protein response (UPR) plus integrated stress response (ISR) pathways triggered by protein misfolding and other stressors, converge on ATF4 as a signalling hub (Fig 1I). We treated female Akita islets with PG *in vitro* for 48h and measured ATF4 as a central readout of stress. ATF4 in Akita or PG-treated islets did not differ from WT (Fig 1J). It is possible that by 4 weeks, acute activation of ATF4 subsides once stress becomes chronic, explaining an absence of ATF4 translation, or that protein at the translational level does not match gene products at the transcriptional level. We measured *xbp1*, *atf4* and *chop* mRNA and did not detect significant differences in WT versus Akita, or PG-treated Akita islets (Fig 1K). Akita mice are characterized by genetically induced insulin misfolding as the primary insult; the development of ER stress, loss of β cell mass, and hyperglycemia are secondary consequences. Given that even modest hyperglycemia can reduce typical maturity transcripts in β cells^35^, we measured identity transcription factors (*pdx1, mafa* and *nkx6.1*) in islets from Akita: only *mafa* was significantly decreased in Akita compared to WT and was not normalized by PG (Fig 1L).

Since the prosurvival and protective effects of PG were unexpectedly detached from ER stress markers, we hypothesized that PG alleviates β cell dysfunction in Akita mice^36^. We noted an increase in functional glucose response via seahorse in PG-treated islets (Fig 1M, Fig 1N). PG-treated Akita islets mirrored WT glucose-responsiveness, particularly in extracellular acidification (ECAR) versus oxygen consumption (OCR), indicating a likely change in glycolytic function and bioenergetics (Fig 1M). Improved β cell function was instead an obvious outcome of PG treatment. The characterization of Akita mice as strictly an ER stress model has obscured the fact that UPR and ISR related genes in Akita islets (i.e. *atf3, atf4, atf6, chop*, etc.) show mixed expression patterns in our data and others^37^. Additionally, Akita islets present with metabolic and bioenergetic defects including reduced oxygen consumption, reduced total glutathione (GSH), oxidative stress and increased mitochondrial ROS that are partially reflective of islet phenotypes in clinical T1D and T2D ^36^. We broadly questioned the types and magnitudes of other stress(es) elicited by protein misfolding in β cells and islets and, more specifically in this study, whether the protective effects of Rnls inhibition during β cell stress arise from metabolic mechanisms that might relate to intracellular antioxidant status.

### Renalase inhibition alters glucose metabolism enzyme expression in β cells

Rnls is a near-ubiquitously expressed intracellular oxidase enzyme with a flavin dinucleotide (FAD) binding domain and structural similarities to the MAO enzyme family^38,39^. Rnls was initially reported to metabolize catecholamines but this was later refuted^40^. The function(s) and subcellular localization of Rnls protein are uncharted. Rnls has predicted enzymatic preference for oxidizing 2-and 6-dihydro NAD(P) isoforms – sometimes annotated as isomeric forms of β-NAD(P)H – that are spontaneously derived from the reduction of β-NAD(P)^41^. The substrates of Rnls are thought to potently inhibit dehydrogenases; based on these *in silico* characterizations, Rnls inhibition could hypothetically mediate stress protective phenotypes by its regulation of glucose metabolism. It is essential to first understand the cellular role(s) of Rnls during homeostatic cellular conditions.

We performed bulk RNA sequencing (RNAseq) in NIT-1 mouse β cells previously generated to harbour CRISPR-induced indel mutation in the Rnls gene (Rnls^mut^)^27^, rendering the Rnls enzyme dysfunctional. Although this approach did not target Rnls expression *per se*, expression of Rnls transcript was decreased (Fig 2A; RNAseq data, left), also via qPCR (Fig 2A, right). Since cell metabolism is often heavily compartmentalized, we measured Rnls localization to gain an appreciation of where it functions within the cell. In β cells, Rnls protein is primarily localized to the cytoplasm, with some expression in the mitochondria (Extended Data 2A).

**Figure 2:**
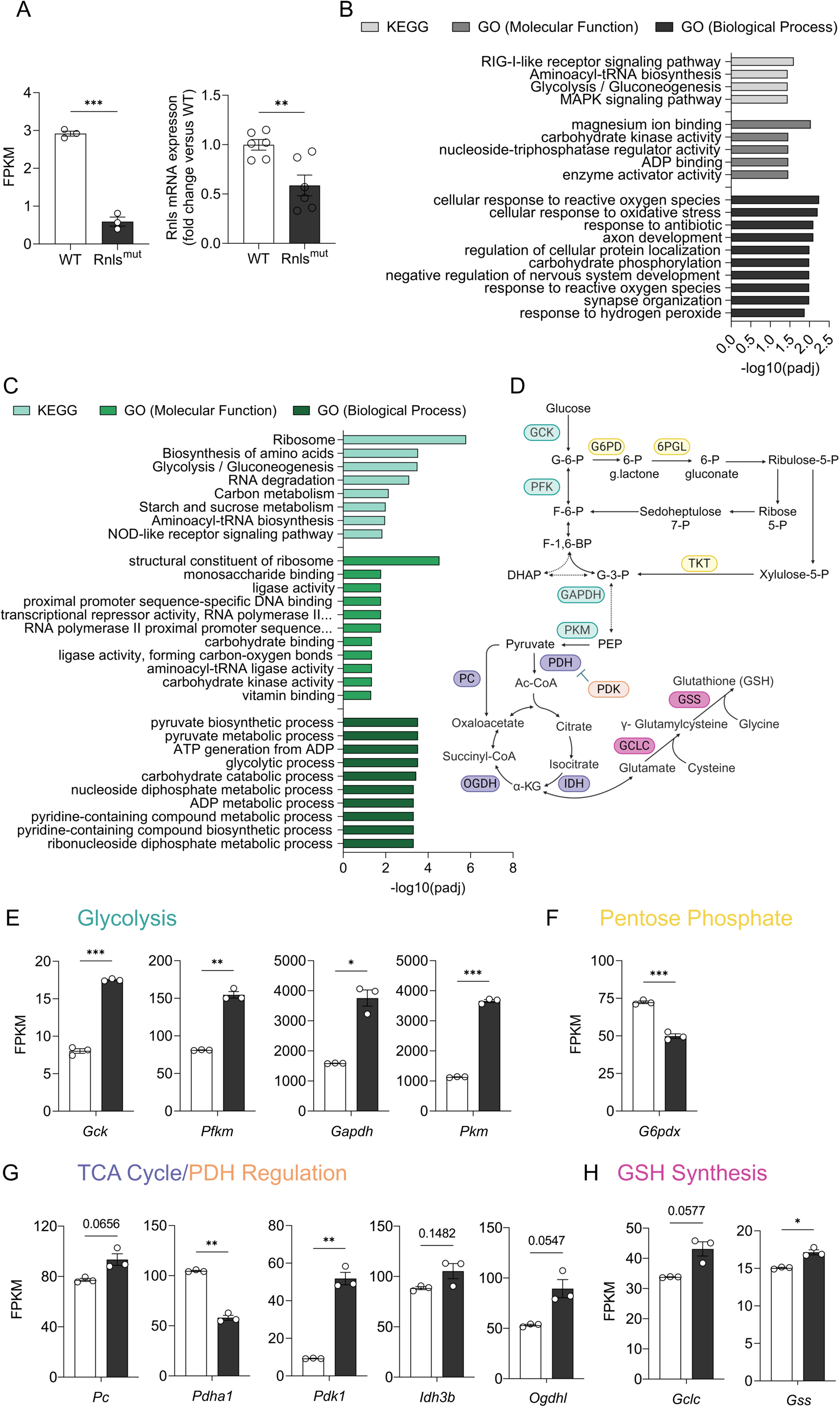
Glucose metabolism and oxidative stress response pathways are regulated by Rnls inhibition β cells. Rnls expression in WT vs Rnls^mut^ NIT-1 β cells measured by (A) RNAseq (left; n=3) or qPCR (right; n=6). (B) Significantly changed KEGG and GO pathway analyses of WT vs Rnls^mut^ NIT-1 β cells (n=3/cell line for all RNAseq analyses) and (C) Significantly increased KEGG and GO pathways in Rnls^mut^ β cells (n=3); all shown are p<0.05 FDR versus WT controls. (D) A schematic diagram of predicted metabolic nodes regulated by Rnls, based on pathways identified in Fig B and C. Rate-limiting enzymes in (E) glycolysis; (F) Pentose Phosphate; (G) PDH regulation and TCA cycle pathways; and (H) GSH synthesis in WT versus Rnls^mut^ NIT-1 β cells. Data is expressed as mean ± SEM. Statistical significance indicated by *p ≤ 0.05; ** p ≤ 0.01; *** p ≤ 0.001 as indicated. Dots represent technical replicates. Abbreviations: FPKM, Fragments Per Kilobase of transcript per Million mapped reads; KEGG, Kyoto Encyclopedia of Genes and Genomes; GO, Gene Ontology; gck/GCK, glucokinase; GSH, glutathione; pfkm/PFK, phosphofructokinase; gapdh/GAPDH, glyceraldehyde-3-phosphate dehydrogenase; pkm/PKM, pyruvate kinase M1/M2; pc/PC, pyruvate carboxylase; pdha1/PDH, pyruvate dehydrogenase (E1 subunit alpha 1); pdk1/PDK, pyruvate dehydrogenase kinase; idh3b/IDH, isocitrate dehydrogenase; ogdhl/OGDH, oxoglutarate dehyodrogenase (analagous to α-ketoglutarate dehydrogenase); G6PD, glucose-6-phosphate dehydrogenase; 6PGL, 6-phosphogluconolactonase; TKT, transketolase; gclc/GCLC, glutamate-cysteine ligase catalytic subunit; gss/GSS, glutathione synthetase.

To determine the biological systems regulated by Rnls inhibition in Rnls^mut^ β cells, we conducted unbiased pathway analyses. The most robust KEGG and GO patterns highlighted glycolysis/gluconeogenesis, carbohydrate phosphorylation and reactive oxygen species/oxidative stress as broadly modified in Rnls^mut^ β cells (Fig 2B). We explored these pathways deeply to assess directionality and discovered that oxidoreductive pathways, glycolysis and pyruvate biosynthesis, carbohydrate catabolism, and amino acid metabolism were upregulated in Rnls^mut^ β cells (Fig 2C). An increase in ribosome pathways in Rnls^mut^ β cells (Fig 2C) was unexpected, given that Rnls is an oxidase enzyme with putative relationships to intracellular NAD(P)H/NAD(P)+ and no known ribosomal annotations. The robust metabolic signature in Rnls^mut^ β cells seemed rational by contrast. Hyperglycemia downregulates translation of insulin mRNA, as well as secretory granule formation, exocytosis and metabolism-coupled secretion^25^. Thus, there is likely a connection between glucose metabolism, ribosomes, and ER stress that has not been explored in the context of Rnls inhibition. Less statistically significant but more widespread increases appeared in carbon, starch, sucrose and carbohydrate metabolism (Fig 2C). The potential points of metabolic regulation and rate-limiting enzymes predicted by RNAseq are shown in Fig 2D.

We next began to reason through likely metabolic mechanisms of stress protection based on the literature. Glucokinase activation is one known mechanism by which glucose metabolism is protective against apoptosis^42^. Downstream, pentose phosphate pathway activity interfaces with glycolysis and antioxidant status as a site of reducing oxidized NADP back to intracellular NADPH, for the purpose of reducing GSH to maintain a favourable redox status, and for other functions^43,44^. However, the contribution of pentose phosphate activity to total oxidative glucose utilization is debatable and has been reported as intriguingly low in β cells^45^. Generating a sufficient pool of NADPH necessary for stress responses in β cells may depend on pyruvate cycling and/or mitochondrial membrane potential, or could be sufficient from pentose phosphate activity despite low flux^46,47^. This is unproven. Linking pentose phosphate mechanisms back to Rnls would require deep investigation into NADP/NADPH dynamics that have yet to be studied. Further downstream, glucose-derived carbon routing through pyruvate carboxylase (PC) is known to enhance the synthesis of major antioxidant metabolite glutathione (GSH) in islets undergoing inflammatory and oxidative stress^48^. There is significant complexity in untangling the many fates of glucose and associated stress protection.

As a first approach, we decided to evaluate the mRNA expression of rate limiting enzymes in each pathway to hypothesize relative pathway importance. In core glucose metabolism, every glycolytic enzyme increased >2x in Rnls^mut^ versus WT β cells (Fig 2E; Extended Data 2B, 2C) alongside a decrease in most TCA cycle transcripts (Fig 2G; Extended Data 2E). Pyruvate dehydrogenase kinase (pdk1), which serves to limit flux through the TCA cycle via PDH, also increased (Fig 2G). In pathways relevant to antioxidant defense status, pentose phosphate enzymes had divergent profiles (Fig 2F; Extended Data 2D), however, biosynthetic GSH enzymes GCLC and GSS were slightly increased in Rnls^mut^ β cells (Fig 2H). Taken together, the largest magnitude of gene expression change in Rnls^mut^ cells was evident in glycolysis itself. Rate-limiting enzymes PFK, GAPDH and PKM were much more highly expressed compared with other pathways and differential expression exceeded patterns in pentose phosphate, TCA cycle or GSH synthesis (Fig 2E versus 2F, 2G, 2H). While transcript-level analyses provide insight into potential mechanisms of Rnls inhibition, a caveat of RNAseq in metabolism is that enzyme gene expression does not faithfully reflect metabolite abundance or metabolic pathway flux and function.

### Glycolytic intermediates, amino acid pools and GSH are increased in Rnls^mut^ β cells

To determine whether Rnls inhibition truly regulates cell metabolism beyond transcriptional reprogramming, we next performed steady-state, untargeted metabolomics to measure steady-state metabolite abundances in WT versus Rnls^mut^ β cells. Pathway enrichment (Extended Data 3A) confirmed that Rnls inhibition in Rnls^mut^ β cells impacts core metabolic pathways including the TCA cycle, multiple amino acid pools, pyruvate-related metabolism and GSH; the latter two signatures aligned strongly with RNAseq patterns. Glucose-derived metabolites and amino acid abundances were next mapped into (simplified) pathways (Fig 3A). The most striking increases in steady-state metabolite pools were evident in glycolytic intermediate 2-PG and amino acid glutamate, serine, glycine and homocysteine pools. These data would suggest elevated glycolysis and an increase in amino acid precursors to GSH synthesis during Rnls inhibition in Rnls^mut^ β cells. Linking glucose-derived GSH to stress protection is conceptually obvious^48,49^. Glycolysis, on the other hand, decreases ∼25% in diabetic islets^50,51^. The impact of correcting glycolytic flux, or regulation of β cell function and viability by glycolytically derived metabolites, reducing equivalents (i.e. NADH) and/or ATP are unknown.

**Figure 3:**
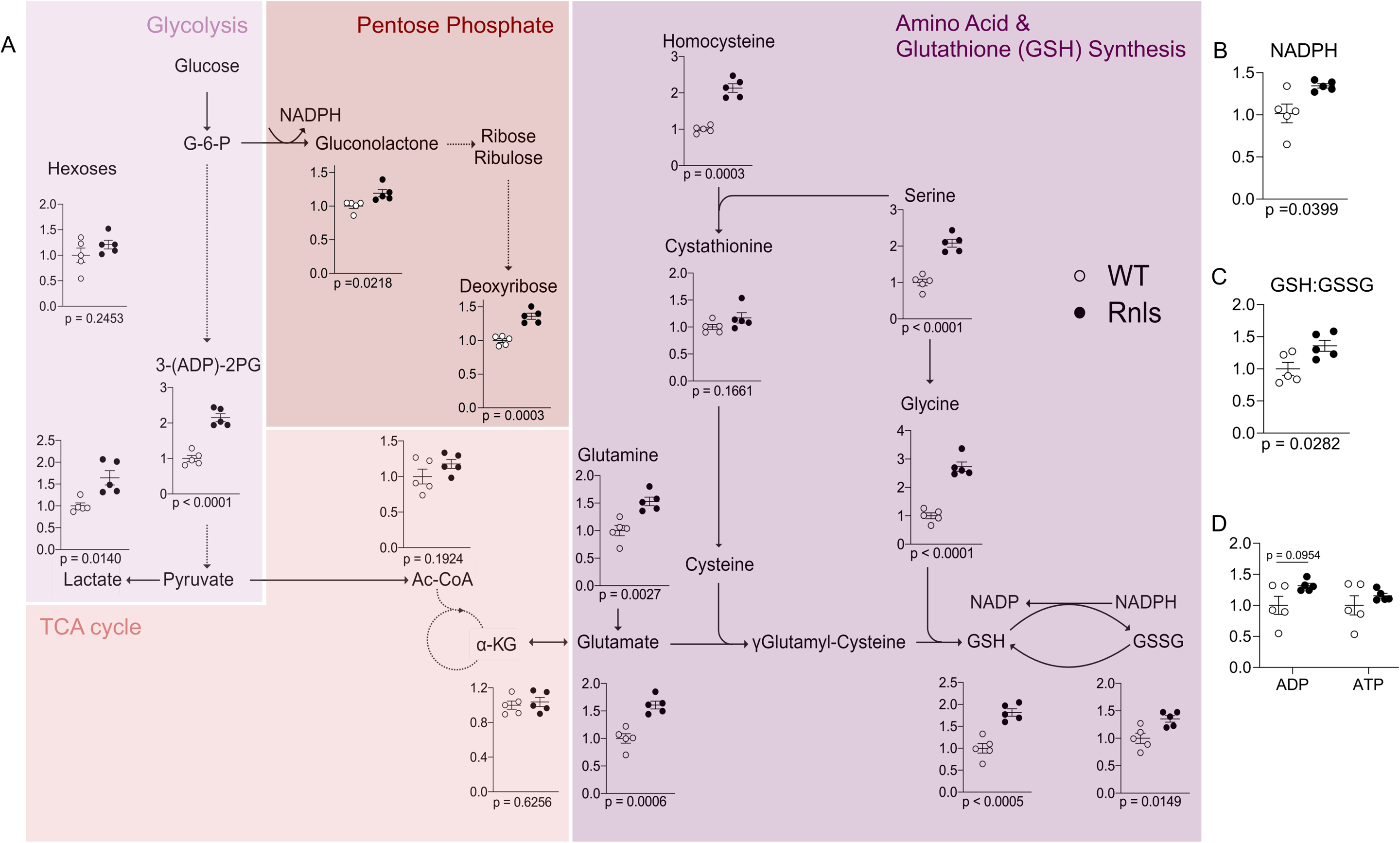
Untargeted metabolomics predicts increased glycolytic intermediates, amino acids and GSH metabolite pools in Rnls^mut^ β cells. WT and Rnls^mut^ NIT-1 β cells were collected in basal conditions to assess metabolite abundances. (A) Abundance of metabolites related to glycolysis, TCA cycle, pentose phosphate and GSH synthesis – identified through Metaboanalyst in Extended Data Fig 3A – normalized to WT. (B) Quantification of NADPH; (C) Ratio of GSH (reduced): GSSG (oxidized); and (D) ADP and ATP in WT versus Rnls^mut^ cells. Data is expressed relative to WT as 1.0, mean ± SEM and p-values for each metabolite are indicated below each plot. Each dot represents a biological replicate, n=5. Solid line arrows represent one reaction; dotted arrows indicate multiple reaction steps. Abbreviations: G-6-P, glucose-6-phosphate; 3-(ADP)-2PG, 3-ADP-2 phosphoglycerate; Ac-CoA, acetyl coenzyme A; α-KG, α-ketoglutarate; GSH, reduced glutathione; GSSG, oxidized glutathione/glutathione disfulfide; NADP, nicotinamide adenine dinucleotide phosphate (oxidized); NADPH, nicotinamide adenine dinucleotide phosphate (reduced).

Apart from glucose-related metabolite pools, the relative availability of ADP, ATP or other co-factors including NAD(P)+ or NAD(P)H could theoretically mobilize stress-protective mechanisms in Rnls^mut^ β cells. NADPH was significantly increased during Rnls inhibition in Rnls^mut^ β cells (Fig 3B). Glutathione reductase (no change in expression; Extended Data 3B) uses NADPH to reduce oxidized glutathione (GSSG) back to GSH, thereby increasing intracellular reductive potential to limit oxidative stress. An increase in NADPH could be derived from pentose phosphate activity, TCA-linked malate or citrate dehydrogenases, anaplerotic pyruvate cycling or potentially via Rnls inhibition directly, although the latter possibility is less likely. The expression of antioxidant enzymes varied in Rnls^mut^ β cells (Extended Data 3B). Changes in antioxidant status during Rnls inhibition are likely due to NADPH, GSH availability and/or elevated GSH:GSSG (oxidized glutathione; Fig 3C) as opposed to differences driven by enzyme expression. Whole-cell ATP and ADP did not differ in Rnls^mut^ cells (Fig 3D), indicating no baseline change in metabolic coupling factors for insulin secretion or high energy phosphate advantage for other biochemical reactions.

Since Rnls also regulates amino acid and GSH metabolite pools, it is entirely possible that posttranslational mechanisms, including glutathionylation ^52^, sumoylation^53^, etc. may also contribute to a bioenergetic basis for β cell survival and/or function during stress.

### Basal and stress-induced cellular bioenergetics are altered during Rnls inhibition in Rnls^mut^ and Pargyline-treated β cells

Both identification of elevated glycolytic intermediates in absence of alterations to the TCA cycle in Rnls^mut^ β cells and an increase in ECAR in Akita mouse islets treated with PG would predict an increase in glycolysis during genetic or pharmacological Rnls inhibition. A change in actual β cell function would provide proof of a *bona fide* metabolic shift during Rnls inhibition. We used Seahorse experiments to address three important questions: whether bioenergetics were altered in Rnls^mut^ β cells; if pharmacological Rnls inhibition works along the same mechanisms as genetic Rnls^mut^ cells; and, critically, if Rnls inhibition impacts the metabolic response to an acute stress challenge. We tested the hypothesis that Rnls inhibition alters glycolysis and/or TCA cycle activity, on the premise that the interaction of stress pathways and cell metabolism must underlie stress-protective responses based on RNAseq data and prior phenotypes^27^. Extracellular acidification rate (ECAR) – a proxy for glycolysis – was elevated basally in Rnls^mut^ β cells (Fig 4A, 4B) to a greater extent than changes in OCR (Extended Data 4A, 4B). This observation is consistent with transcriptional and metabolite profiles hinting towards reliance on glycolysis and confirms a true change in β cell bioenergetic and metabolic function during Rnls inhibition.

**Figure 4:**
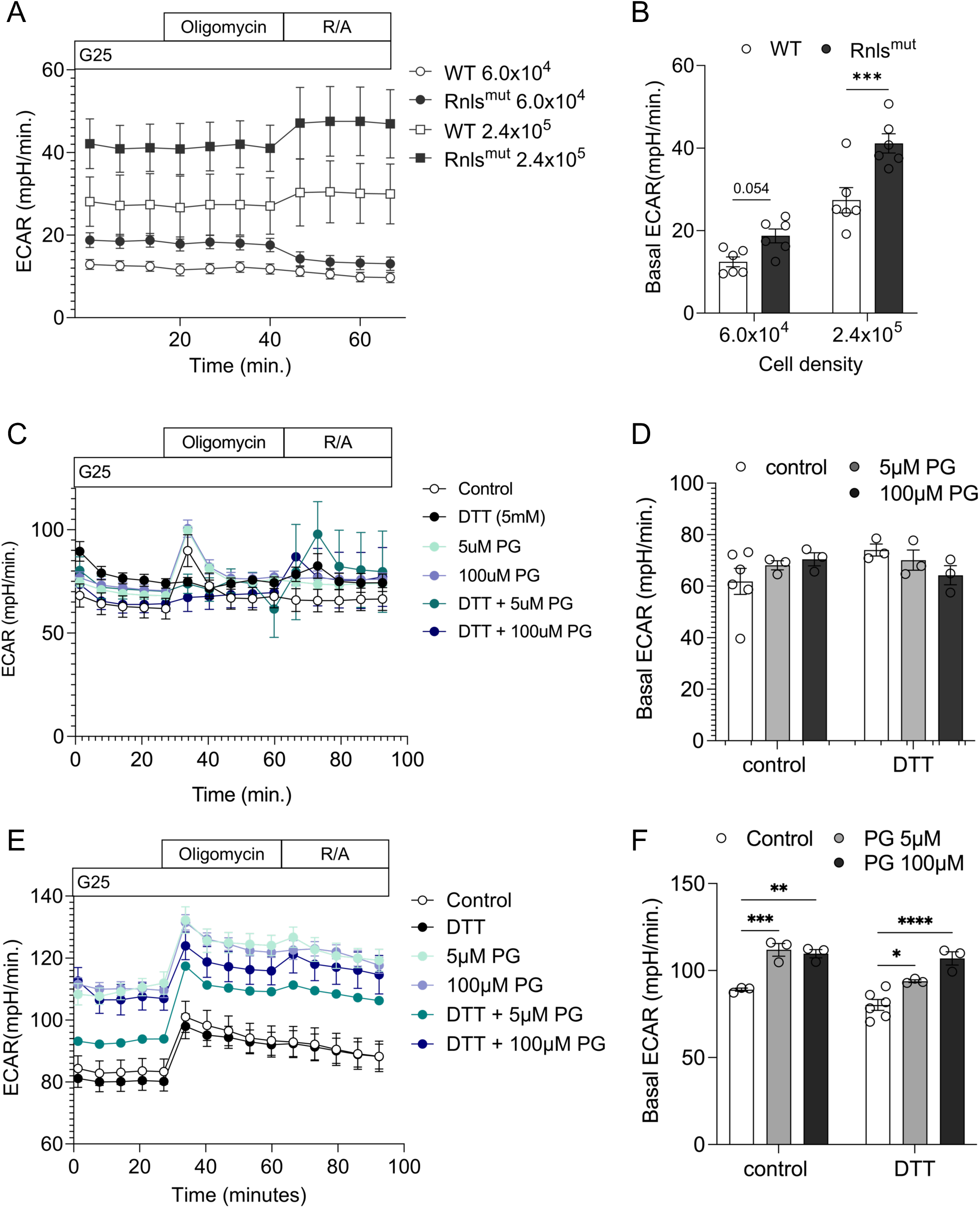
Rnls inhibition regulates basal β cell metabolism and acute bioenergetic stress responses. Extracellular acidification (ECAR), a proxy for glycolysis, measured in (A) WT versus Rnls^mut^ NIT-1 β cells, with basal ECAR quantified in (B). NIT-1 cells treated with 5-100µM Pargyline (PG) for 24h and challenged with DTT for 1h in (C). Basal ECAR is quantified in (D). MIN6 cells treated with 24h PG and challenged with DTT for 1h in (E) and basal ECAR quantified in (F). Data is expressed as mean ± SEM. Statistical significance indicated by *p ≤ 0.05; ** p ≤ 0.01; *** p ≤ 0.001; *** p ≤ 0.0001 as indicated. Data is n=3-6 technical replicates from one representative experiment repeated n=3 times in WT versus Rnls^mut^ NIT-1 β cells; repeated n=2 times in WT vs PG-treated NIT-1; or repeated n=2 times in WT vs PG-treated MIN6.

Our experiment design was expanded to include two concentrations of Pargyline (PG) as a pharmacological comparison to genetic inhibition in Rnls^mut^ cells. Rnls^mut^ was previously shown to protect against acute ER stress triggered by thapsigargin (TG)-induced Ca^2+^ influx^27^. Here we instead aimed to induce protein misfolding *in vitro* in β cells using dithiothreitol (DTT), since insulin misfolding occurs as a true physiological perturbation in stressed islets, plus this stressor complements *in vivo* experiments in Akita mice (see Figure 1)^32,54–56^. In NIT-1 cells treated with PG, basal glycolysis increased 20%. This magnitude was less significant than the change in Rnls^mut^ β cells (Fig 4C, 4D; Extended Data 4C, 4D). We questioned whether ECAR in the WT NIT-1 model was too close to measurement sensitivity and extended experiments to test NIT-1 bioenergetic profiles against another insulinoma line, MIN6 β cells, in the presence of PG and at 5mM low glucose DMEM to push cells away from high reliance on glycolysis. Culturing insulinoma cells in low glucose media did not substantially change their bioenergetic characteristics (Extended Data 4E). However, MIN6 were considerably less glycolytic than NIT-1 β cells at baseline, as confirmed by increased mitochondrial ATP production and less sensitivity to 2-deoxy-D-glucose (2-DG; Extended Data 4E). Pharmacological Rnls inhibition with PG increased baseline ECAR in MIN6 to mimic Rnls^mut^ cells (Fig 4F). This bioenergetic shift was sustained during ER stress challenge with DTT and suggests that Rnls inhibition changes metabolic and bioenergetic responses to acute stress (Fig 4E, 4F). OCR did not change at rest in the same 5µM PG-treated MIN6 cells (Extended Data 4F, 4G).

A bioenergetic increase in ECAR is likely influenced by lactate, which was increased in Rnls^mut^ β cells (Fig 3A) and might be considered an artefact of insulinoma cells. Conversion of pyruvate to lactate is typically thought to be repressed in β cells. However, a recent shift in our canonical understanding of β cell/islet lactate dehydrogenase (LDH) suggests that lactate is indeed produced by LDH in islets, especially in human donors, with unknown impacts on β cell function and viability, and that this pathway may hold new relevance in primary β cells and islets^57^. The role of Rnls in LDH-mediate lactate production is unknown.

Overall, genetic mutation in Rnls^mut^ or pharmacological PG-mediated Rnls inhibition in β cells are both consistent with a basal metabolic switch towards glycolysis that is sustained during acute stress. More broadly, acute stress-induced metabolic and bioenergetic changes are consistent with enrichment of glycolytic intermediates and increased ECAR in chronically ER-stressed MIN6 cells^58^. It was not our primary aim to investigate broader metabolic responses and adaptations to stress in β cells, but our data do raise a crucial question: in other settings, rapid ATP production and biosynthesis via aerobic glycolysis are advantageous^59^; is this also the case in “stressed” β cells or islets? Seahorse is best used for broad phenotypic screening given its known caveats^60^. ECAR cannot be purely ascribed to glycolysis unless paired with metabolomics to confirm pathway activities, since CO_2_ released from mitochondrial oxidative pathways also contributes to acidification. Oxygen consumption is also not strictly mitochondrial and is the summation of other O_2_-dependent intracellular biochemistry. We followed bioenergetics assays with independent [^13^C_6_]-glucose tracer experiments to confirm whether Rnls-regulated bioenergetics coincides with true changes in cell metabolism.

### Glycolysis and GSH synthesis are augmented during acute ER stress in β cells during Rnls inhibition

We used data from Figure 2 to guide targeted metabolic tracer experiments in the context of β cell stress. Given that Rnls^mut^ showed phenotypes of increased glycolysis, augmented amino acid pools and GSH enrichment at both transcript and metabolite levels, we used [^13^C_6_]-glucose isotope tracer to measure the steady-state flow of glucose-derived carbons during acute stress, with the expectation of increased labelling in glycolytic intermediates and carbon shunting from the TCA cycle towards GSH synthesis. To mimic metabolic function experiments, ER stress was induced using 1h DTT to activate the UPR and ISR protein ATF4 without causing β cell death (Extended Data 5A).

Potential [^13^C_6_]-glucose-derived carbon labelling from glycolysis to the TCA cycle through GSH synthesis is depicted in Fig 5A. We noted a considerable increase in [^13^C_6_]-glucose incorporation into the glucose-6-phosphate (G-6-P) pool basally and especially in response to 1h DTT exposure in Rnls-inhibited β cells (Fig 5B). This finding is again congruent with augmented glycolysis during Rnls inhibition basally and in response to cellular stress. Very rapid sampling is required to capture full glycolytic dynamics (i.e. 10s), and slightly less strict for TCA cycle metabolites (>2h), therefore distal glycolytic intermediates are not shown. Nonetheless, this pattern of G-6-P labelling is consistent with elevated 2-PG abundance and increased ECAR in Rnls inhibited β cells identified in Figures 3 and 4. Interestingly, PG phenocopied G-6-P labelling in Rnls^mut^ cells to confirm that pharmacological inhibition at least partially mimics metabolic mechanisms discovered in the genetic Rnls^mut^ cell model.

**Figure 5:**
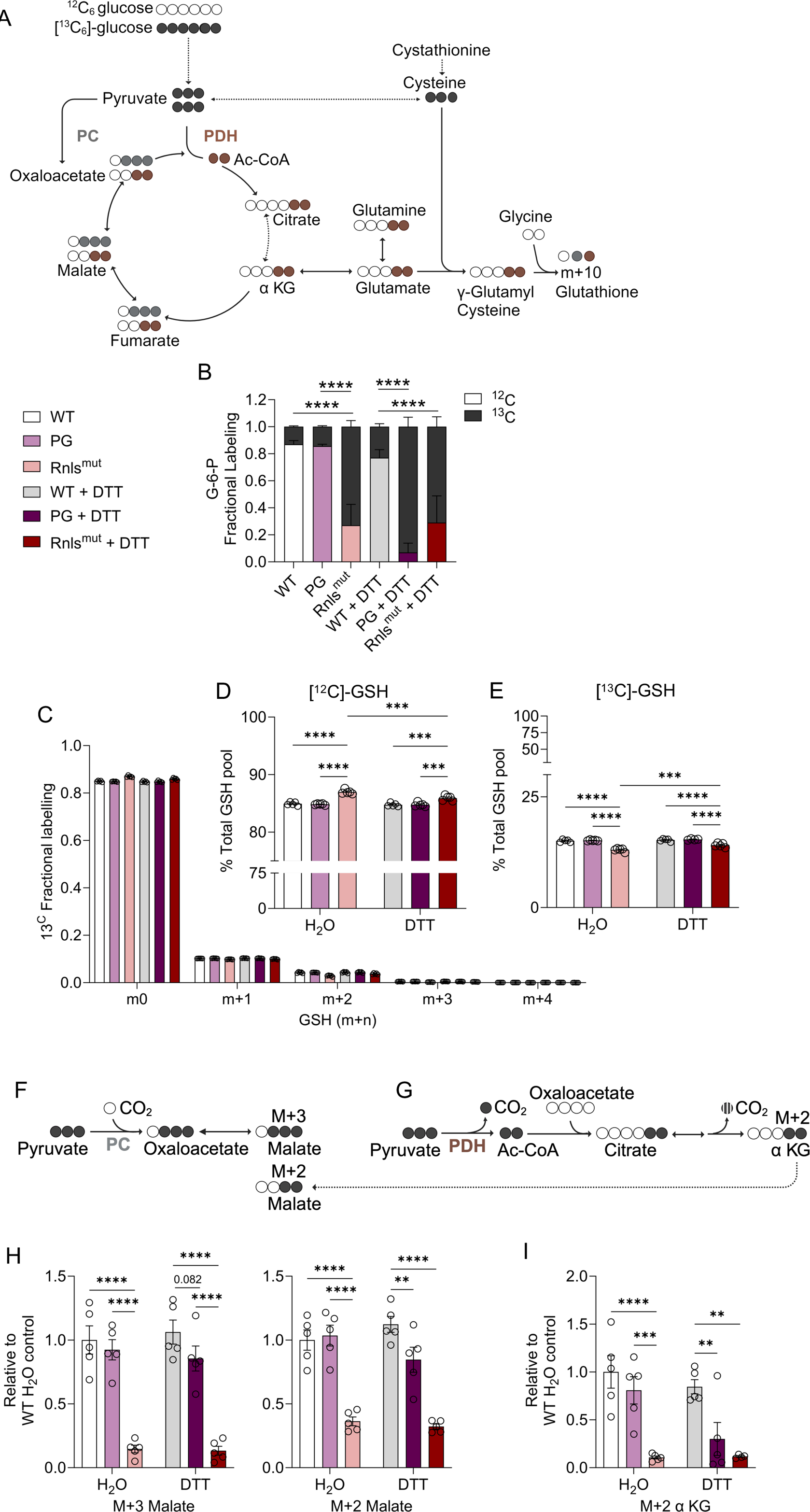
Rnls inhibition promotes a metabolic shift to glycolysis and GSH synthesis during acute stress. Wildtype (WT) control, Pargyline (PG)-treated and Rnls^mut^ (Rnls) NIT-1 β cells were cultured in media containing [^13^C_6_]-glucose for 24h and challenged with 1h DTT stress to stimulate acute stress via protein misfolding. (A) Potential labelling routes for [^13^C_6_]-glucose incorporation through glycolysis, the TCA cycle and GSH synthesis. Labelling via anaplerotic pyruvate carboxylase (PC) flux is shown in gray, and oxidative pyruvate dehydrogenase (PDH) flux in brown. (B) Relative ^13^C_6_ labelling of the first glycolytic intermediate, glucose-6-phosphate (G-6-P), normalized to WT control. Downstream [^13^C_6_]-glucose incorporation into GSH is shown in (C): total fractional labelling for m+0 (unlabelled) through m+n (labelled) GSH istopologs, up to m+4; m+5 through m+10 isotopologs did not accumulate. Relative incorporation of glucose into (D) unlabelled [^12^C_6_]-GSH, versus (E) labelled [^13^C_6_]-GSH from the total sum of the intracellular GSH pool. (F-G) Schematic showing the potential fates of m+3 pyruvate containing 3 carbons derived from [^13^C_6_]-glucose through PC and PDH in one turn of the TCA cycle. (F) Pyruvate is irreversibly carboxylated to oxaloacetate through PC to generate m+3 oxaloacetate, which is reduced to m+3 malate. (G) Oxidative decarboxylation of m+3 pyruvate to m+2 acetyl-coA via PDH (one carbon is lost via CO_2_). Citrate synthase metabolizes oxaloacetate and acetyl-coA to generate m+2 labelled citrate (followed by isocitrate; not shown), m+2 α-ketoglutarate and m+2 malate in traditional TCA cycle flux. (H) Labelled m+3 and m+2 malate relative and (I) m+2 α-ketoglutarate relative to WT control. Data is n=5 biological replicates. Data is expressed as mean ± SEM. Statistical significance indicated by ** p ≤ 0.01; *** p ≤ 0.001; *** p ≤ 0.0001 as indicated.

Since Rnls^mut^ β cells were characterized by an increased GSH:GSSG ratio, we measured steady-state GSH synthesis during Rnls inhibition and changes in GSH biosynthesis during acute ER stress. Total fractions of m+0 (unlabelled) through m+n ([^13^C_6_]-glucose labelled) GSH are consistent with experiments in islets that demonstrate ∼20-25% labelling over 24hrs^48^. Genetic Rnls inhibition in Rnls^mut^ β cells increased incorporation into ^13^C-GSH during DTT stress (Fig 5E) alongside a concomitant decrease in unlabelled ^12^C GSH, as expected (Fig 5D); this change in GSH synthesis was slight but remarkable given that the period of stress exposure was only 1h. This pattern was not phenocopied by PG treatment and suggests that Rnls^mut^ more potently regulates metabolic mechanisms and phenotypes, consistent with stronger phenotypes also noted in Seahorse data (see Figure 4). Partial specificity of PG for Rnls could also explain a weaker metabolic effect in both seahorse and metabolomics analyses. It is also possible that a more chronic stress exposure would be required to detect PG-mediated changes in distal GSH incorporation. Taken together, glucose-derived GSH labelling during acute ER stress in Rnls^mut^ β cells contributes to a more favourable intracellular redox profile by increasing the GSH:GSSG ratio alongside augmented NADPH (Fig 3B). Given the discordant changes in antioxidant enzyme expression in Rnls^mut^ β cells (Extended Data 3B), incorporation of glucose into GSH during stress more likely contributes to β cell protection than transcriptional changes in GSH and redox enzymes. These data imply a direct connection between Rnls activity, ER stress responses, and at least two new mechanisms that regulate β cell metabolism during Rnls inhibition: an increase in glycolysis and a boost in GSH antioxidant defense capacity.

Seahorse and untargeted metabolomics experiments hinted at minor changes in mitochondrial pathways concomitant to altered glycolysis. Presumed Rnls substrates 2 and 6-dihydroNAD(P) have high affinity for *E.coli* malate dehydrogenase (MDH) to inhibit its activity^41^. In theory, Rnls activity would alleviate a “brake” by metabolizing 2 and 6-dihydroNAD(P), and conversely, its inhibition would be predicted to limit MDH activity and decrease intracellular malate. Our data support this model by showing decreased m+2 and m+3 malate generation in Rnls^mut^ and PG-treated β cells. Interestingly 2- and 6-dihydroNAD(P) do not engage with core TCA cycle enzymes PDH or the α ketoglutarate decarboxylase complex, suggesting isoform-specific regulation of dehydrogenases by Rnls that requires further elucidation. It is also not clear whether the deviation towards glycolysis during Rnls inhibition is directly derived from transcriptional upregulation of glycolytic enzymes, or indirectly a consequence of dehydrogenase inhibition and reduced TCA cycle flux. [^13^C_6_]-glucose routing of glucose-derived glycolytic metabolites through anaplerotic PC (Fig 5F) or oxidative PDH (Fig 5G) into the TCA cycle could theoretically be disrupted by Rnls-mediated shift in bioenergetics towards glycolysis. We observed decreased m+3 and m+2 malate (Fig 5H), in basal Rnls^mut^ β cells as well as during ER stress in PG-treated cells. This pattern was consistent in m+2 α-ketoglutarate (Fig 5I) and supports the conclusion that Rnls inhibition favours glycolysis and siphons carbons away from the TCA cycle towards GSH synthesis.

In previous work we found that Rnls^mut^ cells were better-equipped to defend against acute (5h) thapsigargin (TG) ER stress and hydrogen peroxide (H_2_O_2_) oxidative stress. Here we questioned whether PG might maintain NIT-1 cell viability during a more chronic (>24h) stress exposure, versus 1h acute DTT stress, and whether pharmacological Rnls inhibition would protect against TG and H_2_O_2_. Indeed, PG defended wildtype β cells against DTT, TG and H_2_O_2_ (Extended Data 5D-F). These data suggest that metabolic alterations imparted by Rnls inhibition translate to real, broadly pro-survival effects in β cells on subchronic timescales.

### The metabolic basis of Rnls-induced stress protection is conserved in human stem cell-derived β cells

There is great promise in generating sc-β cells for transplantation in T1D. However, these cells are susceptible to immune destruction and remain functionally immature. A “bottleneck” in sc-β cell function is attributed to an issue in glycolysis at the level of GAPDH^61^. We surmised that Rnls inhibition in human sc-β cells might correct metabolic defects to improve functionality by increasing glycolytic machinery at the transcriptional level, as observed in Rnls^mut^ NIT-1 β cells, by mirroring increased glycolysis noted in NIT-1 and MIN6 insulinoma lines. To test whether Rnls inhibition alters glycolysis, TCA cycle, or GSH pathways to impact diabetogenic stress outcomes in human β cells, we differentiated both WT and clonal Rnls knockout (Rnls KO; Fig 6A) human pluripotent stem cells (iPSCs) to sc-β cells. iPSCs were obtained from the Melton lab and maintenance carried out as described, and Rnls KO cells were generated by CRISPR–Cas9 gene targeting^27^. Consistent with previous data, at stage 6 the differentiated iPSCs expressed ∼20-30% c-peptide+/NKX6.1+ co-positive sc-β cells (Fig 6B) and both lines formed intact islet-like clusters upon aggregation (sc-islets; Extended Data 6B) with no appreciable differences in quality or differentiation efficiency (Extended Data 6A).

**Figure 6:**
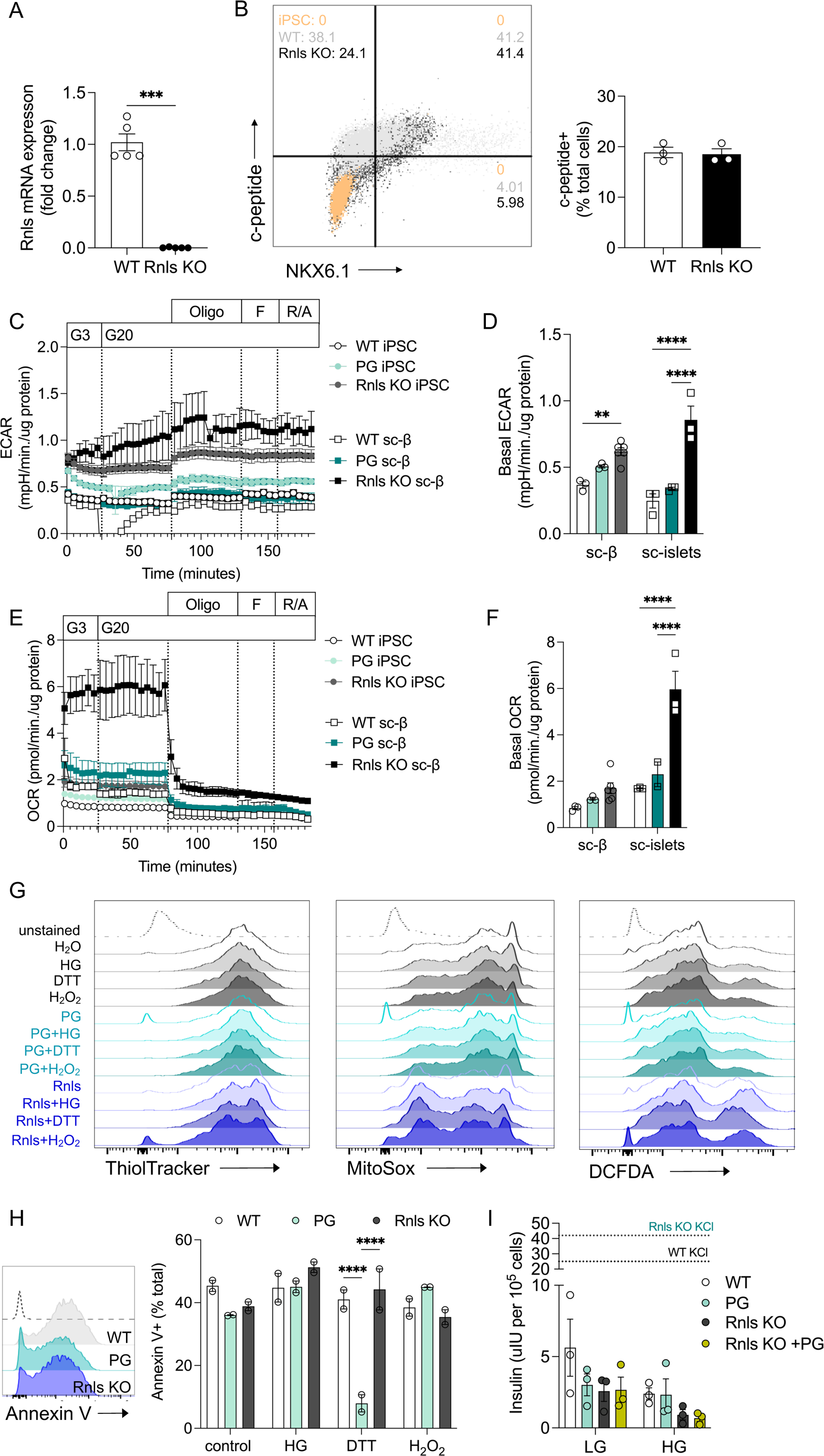
Rnls KO shifts human stem cell-derived β cells towards increased antioxidant capacity and glycolysis. WT and Rnls KO pluripotent iPSCs were differentiated into mature stem-cell derived β cells (sc-β cells). (A) Rnls expression in WT vs Rnls KO iPSC (stage 3). (B) Representative flow cytometry of undifferentiated iPSC (negative control), WT and Rnls KO c-peptide+, NKX6.1+ and double-positive gates. Quantification of total c-peptide+ cells from n=3 differentiations shown; measured by IHC. Planar (sc-β) and aggregate (sc-islets) metabolic function was measured via seahorse in WT, PG (5µM, 24h) and Rnls KO cells. (C) Extracellular acidification rate (ECAR) and (D) basal ECAR in sc-β cells (planar) and sc-islets (aggregates). (E) Oxygen consumption rate (OCR) and (F) basal OCR in the same experiment; n=3-6 replicates shown from one experiment repeated n=3 times. (G) WT, Rnls KO and PG-treated sc-islets were challenged with high glucose (HG), ER stress (DTT) or oxidative stress (H_2_O_2_) for 48h. Histograms show mean fluorescence intensity (MFI) of ThiolTracker, MitoSox and DCFDA. (H) Histogram for Annexin V MFI and quantification of Annexin V+ (apoptotic) cells. (I) Glucose-stimulated insulin secretion (GSIS) in planar sc-β cells in response to low glucose (LG; 2mM glucose) or high glucose (HG; 20mM glucose). Data are expressed as mean ± SEM for n=2-3 differentiations. Each dot represents a biological replicate from n=2-6 technical replicates. Statistical significance *p ≤ 0.05; ** p ≤ 0.01; *** p ≤ 0.001; *** p ≤ 0.0001 as indicated. Abbreviations: G3, 3mM glucose; G20, 20mM glucose; Oligo, oligomycin; R/A, rotenone/antimycin A; DCFDA, 2’,7’-dichlorodihydrofluorescein diacetate.

We questioned whether a pattern of metabolic reprogramming induced by Rnls KO or PG would be mirrored by a shift in bioenergetics towards extracellular acidification in sc-β cells. Rnls KO increased both ECAR (Fig 6C, 6D) and OCR (Fig 6E, 6F) in clustered sc-islets but only ECAR in planar sc-β cells. Consistent with NIT-1 and MIN6 models, glycolysis was increased more substantially in sc-β cells during Rnls inhibition, while oxygen-consuming respiratory pathways remained unaltered or also slightly increased. This is not unusual: increased aerobic glycolysis does not necessitate a suppression in oxidative phosphorylation^62^. PG showed only a partial ability to phenocopy Rnls KO, consistent with our observations that genetic Rnls inhibition is more potent and specific than pharmacological targeting of Rnls. Despite Rnls inhibition increasing rate-limiting glycolytic enzymes and boosting glycolysis, neither PG or Rnls KO corrected the well-documented poor responsiveness of sc-β cells to 20mM glucose (Fig 6C, Fig 6E, 6I). Although Rnls inhibition did increase basal glycolysis in human sc-β cells, the well-documented phenotypes pf poor glucose responsiveness and absent GSIS in sc-β cells persisted in both WT and Rnls KO ^61,63^. One theory is that restricted TCA metabolite labelling and mitochondrial metabolic activity could arise from this aberration in distal glycolysis^64^, however, augmenting glycolysis did not seem to benefit functional outcomes in our sc-β cells. Our data also confirm intact mitochondrial machinery^63^; however, respiratory insensitivity to oligomycin and FCCP in sc-β cells, uncharacteristic of mature human islets, suggest aberrations in coupling to ATP provision that may affect viability due to altered bioenergetics. There was also no change in glucose-stimulated insulin secretion (GSIS) in planar sc-β cells with Rnls KO or PG exposure (Fig 6I). Sc-β cells from both Rnls KO and WT lines were responsive to KCl, suggesting that sc-β cells can depolarize but do not mimic metabolic coupling that is characteristic of mature islets. Nonetheless, metabolic phenotypes are remarkably consistent from mouse to human β cells during Rnls inhibition.

To assess the redox environment in human sc-β cells, we used ThiolTracker‒a proxy for intracellular GSH – as an orthogonal method to metabolomics, to assess whether antioxidant status was also more favourable in Rnls KO and PG-treated human sc-β cells. The mean fluorescence intensity of GSH intensity did not differ in PG treatment or Rnls KO during 24h challenge with high glucose (HG), DTT, or H_2_O_2_ to impose acute metabolic, ER and oxidative stress, respectively (Fig 5G; Extended Data 5Ci). However, a bimodal distribution was visible in the Rnls groups and may indicate a population of cells with high GSH availability; worth noting is that this mature population is still only 20-30% “true” β-like cells, which might affect relative proportions of redox and stress markers. Since the ThiolTracker probe is reflective of both synthesis and turnover of free thiols, we questioned whether GSH in PG and Rnls KO might still be sufficient to buffer intracellular ROS generated during diabetogenic stress. Mitochondrial ROS measured with mitoSOX underwent a significant leftward shift in basal and stress conditions, in both PG and Rnls treatment (Fig 6G; Extended Data 6Cii), indicating reduced mitochondrial oxidative stress. Cytosolic ROS was not regulated by Rnls (Fig 6G; Extended Data 6Ciii). Therefore, antioxidant capacity during Rnls inhibition in human sc-β cells mitigates ROS in the mitochondrial subcellular compartment. Even with retained aberrations in glucose processing in the human sc-β cells, the patterns of augmented antioxidant capacity and altered glucose metabolism seen in mouse NIT-1 were conserved in human β cells with Rnls KO or PG treatment.

We moved to challenging WT, PG-treated and Rnls KO sc-β cells with more chronic stress (24h) and measured apoptosis (Annexin V+; Fig 6H). Rnls inhibition did not significantly impact apoptosis, apart from PG protection against DTT, which we cannot currently explain. Interestingly, this aligns with another report of negligible apoptosis under single-factor metabolic, ER or inflammatory stress in sc-β cells^26^. A combination of stresses, which likely better mimics the natural T1D and T2D islet microenvironment, appears necessary to inflict meaningful sc-β cell apoptosis and diminished viability.

## Discussion

Associations between Rnls and T1D risk and age of onset have appeared in GWAS studies^28,29^. Despite strong evidence for its role in autoimmunity, ER and oxidative stress responses in the context of T1D and sc-β cell transplantation, the function(s) of Rnls and its inhibition remained entirely unknown after its annotation to catecholamine metabolism was deemed untrue. We used a powerful and deductive combination of RNAseq, untargeted and [^13^C_6_]-glucose metabolomics, and bioenergetic phenotyping to illuminate that *in vitro* Rnls inhibition regulates glycolysis and GSH pathways as dual mechanisms underpinning phenotypes of β cell stress protection. To our knowledge, this is the first mechanistic evidence detailing the molecular effects of Rnls (in)activity in mammalian cells and tissues.

Here we advanced fundamental knowledge of Rnls biochemistry by pinpointing metabolic nodes under control of this intracellular oxidase: increased basal glycolysis and amino acid pools, in addition to a favourable antioxidant environment (i.e. elevated GSH:GSSG and NADPH) were robustly conserved from mouse to human β cells during Rnls inhibition, across transcript and mechanistic levels. Glucose metabolism itself can stimulate β cell replication to maintain β cell mass, and non-allosteric pharmacologic activation of glucokinase mitigates apoptosis in β cells and human islets exposed to hyperglycemia and inflammation ^42,48,65–69^. Critical metabolic axes within molecular control of glucokinase ^42,66,70^ – PC as the primary example – achieve β cell stress protection via multiple synergies between interrelated arginine-ureagenesis and GSH synthetic metabolic pathways^48^. Rnls is now emerging with similar complex metabolic reprogramming, likely secondary to changes in isomers of NAD(P)H that are near-impossible to detect using LC/MS^41^. The pro-survival effects of Rnls are attributed to stress-protective mechanisms imparted by proximal (glycolysis) and/or distal (GSH) glucose metabolism. Whether glycolytic intermediates, reducing equivalents or ATP production are independently protective against β cell dysfunction and apoptosis is unknown in contrast to the well-documented redox benefits of GSH and NADPH pools. The question that now arises is which pathway is more stress-protective, augmented glycolysis or NADPH and GSH pools? Work is ongoing to investigate pathways not directly assessed here, including [4-^2^H] and [1-^2^H]-glucose for NADPH generation by pentose phosphate activity versus anaplerotic cycling^71^, plus shorter [^13^C_6_]-glucose timeframes, in order to resolve the relative potency of metabolically-derived stress protective mechanisms during Rnls inhibition.

Although this was not a primary question, we substantiated the predicted role for Rnls in dehydrogenase biochemistry by demonstrating reduced m+2 and m+3 malate in Rnls^mut^ and PG-treated β cells. In *e.coli*, putative Rnls substrates 2 and 6-dihydro NAD(P) potently inhibit MDH, though curiously, do not interact uniformly with all dehydrogenases^41^. We show that in Rnls^mut^ cells with an expected accumulation of these substrates, this biochemistry is conserved in mammalian cells and MDH activity is reduced.

A major pattern in our data is a dominance of genetic Rnls^mut^ phenotypes and molecular mechanisms over pharmacological PG treatment, especially in human sc-β cells. Pargyline is an irreversible inhibitor with only partial selectivity for Rnls, as it also interacts with MAO-B in a putative FAD-binding domain^72^. There are two important conclusions from our observations: further research is warranted to optimize potent small molecule inhibitors against Rnls, without off-target effects on MAOs; and the novel drug mechanisms we present here, stemming from only partial Rnls inhibition, may yield new drug development or identify potential drug repurposing opportunities in developing β cell therapeutics against T1D and/or T2D.

The concept of β cells as perpetrators instead of “victims” of T1D was first proposed in 1985 and hypothesized that β cell suicide contributes to diabetes alongside immune cell homicide ^73^. This framework posits that environmental or intrinsic defect(s) arise to cause β cell dysfunction and/or provoke autoimmunity. In support of this concept, immunotherapies against T1D ‒ anti-CD3+ antibody teplizumab as a remarkable example^74,75^‒show only short-term efficacy in delaying disease^74^. It is now becoming accepted that the autoimmune nature of T1D may be initiated, perpetuated or accompanied by peripheral triggers in the pancreas itself that more typically resemble T2D^18^. In recent years, β cell stress has emerged as a strong candidate for β cell suicide with known contributions to death and dysfunction. Acute stress-induced shifts in β cell metabolism and bioenergetics – at both phenotypic and molecular levels – have largely not been elucidated, despite the understanding of nutrient signaling and metabolic coupling factors in basal β cell glucose metabolism^71,76^. Our data show an intriguing increase in glycolysis during Rnls inhibition in response to acute DTT-induced ER stress in β cells. And yet, this metabolic phenotype was not evident in chronically stressed Akita islets. Therefore, stress-induced metabolic switches may provide bioenergetic advantages that favour β cell survival initially, with unknown consequences on secretion and function that shift to (mal)adaptations once timescales become chronic.

On a chronic timescale, Rnls inhibition via oral PG reduced hyperglycemia, maintained β cell mass, increased β cell maturity markers, and improved islet bioenergetic function in female Akita mice. We sought to determine the molecular mechanisms. In response to misfolded protein, activation of the UPR is essential for resolving acute stress and restoring homeostasis. Chronic UPR signalling along the IRE1ɑ/XBP1 and PERK/ATF4/CHOP axes initiates a pro-apoptotic program before the onset of T1D^19,77^; conversely deleting proximal ER stress effector genes *atf6* and *ire1ɑ* from β cells protects against T1D in mice ^78,79^. We were perplexed by the simultaneous lack of ATF4 protein and *xbp1*, *atf4* and *chop* mRNA in Akita mice. This could be reflective of another limitation: one cannot measure stress in β cells that have succumbed to apoptosis (i.e. dead or dying). The dynamic nature of β cell expansion, β cell loss, and oscillations in stress programming throughout neonatal to adult periods of development suggests that “stress” is not linear, and the ability of PG to stabilize islet function and viability may differ pending timing ^37,80^. Akita mice also present with hyperglycemia secondary to genetically induced protein misfolding and insulin insufficiency. The extent to which hyperglycemia and ER stress participate in a vicious cycle towards β cell dysfunction and loss is not clear. In our hands, modest hyperglycemia was evident by weaning and likely contributed to decreased β cell identity genes *pdx1*, *mafa* and *nkx6.1* in Akita islets^35^. Although PG did not affect the expression of ER stress markers, partial restoration of β cell maturity was an unexpected outcome of PG treatment.

Overall, observations in complementary genetic (Rnls^mut^/Rnls KO) and pharmacologically (PG) inhibited mouse and human β cells illuminated some aspects of Rnls biochemistry and function, basally and during stress conditions. Rnls-regulated prosurvival phenotypes are based on at least two multipronged mechanisms involving glucose metabolism: a bonafide metabolic shift towards glycolysis, and siphoning of glucose-derived carbons away from the TCA cycle towards amino acid pools and GSH synthesis. Beyond its effects on acute stress responses, oral PG treatment in chronically stressed Akita mice led to improvements in hyperglycemia, β cell mass, circulating insulin and islet bioenergetic function. Rnls-derived molecular mechanisms extend beyond cell-intrinsic protection to also influence CD4+ T cell signatures and support the notion that cell therapies may be optimized to work in concert with immunotherapies for T1D^18,81^. Given the ubiquitous importance of glucose metabolism in cells and tissues, and the many etiologies driven by stress, the role of Rnls could evolve to impact other disease areas beyond T1D and T2D.

## Limitations

We rigorously tested patterns garnered *in vitro* by the Akita model *in vivo*. Akita mice are not fully reflective of T1D, as they lack typical immune components, but this model is the best available for testing hypotheses regarding chronic stress and β cell mass. Our analyses were strengthened by including both males and females. We circumvented restrictions in using insulinoma β cell lines by differentiating human sc-β cells to measure key outcomes, in an effort to generate translational, disease-relevant conclusions while studying basic biology.

## Methods

### Experimental models

#### Mice

##### Housing

All experimental procedures were approved and performed in accordance with the Joslin Institutional Animal Care and Use Committee (IACUC) regulations. Mice were housed in pathogen-free facilities at the Joslin Diabetes Center on a 12:12h light:dark cycle and were fed a standard chow diet ad libitum (Lab Diet cat. 5020). Heterozygous male C57BL/6-Ins2^Akita^/J (Akita; strain #: 003548) mice were purchased from the Jackson Laboratory and bred with female C57BL/6 wildtype (WT; strain #: 000664) mice once both reached 8 weeks of age. Littermate controls were used for all experiments. Breeder pairs were retired after 4 litters.

##### Weaning and genotyping

At day 18-19, mice were weaned and genotyped according to the restriction digest assay listed in The Jackson Laboratory protocols. Briefly, tail samples were collected and digested with lysis buffer (50mM KCl, 10mM Tris-HCl, 0,1% Triton-X) and Proteinase K at 56C for <2hrs followed by 95C for 10 mins, and amplified using PCR (Azura cat. AZ-1910). The PCR product was incubated with restriction enzyme Fnu4HI (NEB cat. R0178) for >4hrs at 37C and run on a 3% gel. Akita heterozygotes have bands at 140 and 280bp; WT mice at 140bp.

#### Cell lines

##### NIT-1 β cell culture

Murine NIT-1 β cells (ATCC cat. CRL-2055) were maintained in 25mM high glucose DMEM supplemented with 2 mM L-glutamine (Gibco cat. 25-030-081),10% FBS (R&D cat. 50-152-7066), 50 μM β-Mercaptoethanol (Sigma-Aldrich cat. 60-24-2), and 100 units/mL penicillin/100µg/mL streptomycin (P/S; ThermoFisher cat. 15140122) at 37°C, 5% CO_2_. Rnls^mut^ NIT-1 cells were generated and validated previously^5^. In brief, Rnls gRNA (5′-CTACTCCTCTCGCTATGCTC-3′, MGLibA_46009) was cloned into lentiCRISPR v.2 vector and lentivirus was used to establish cell lines. β-Mercaptoethanol was omitted from 48h stress challenges with DTT, TG, or H_2_O_2_ in NIT-1 because of its capacity to eliminate oxygen free radicals.

##### MIN6 β cell culture

Low passage murine MIN6 β cells were received from Dr. Jun-ichi Miyazaki (Osaka University Medical School) and maintained in 25mM glucose DMEM supplemented with 15% FBS, 0.1mM β-Mercaptoethanol and P/S at 37°C, 5% CO_2_.

#### Mouse islets

##### Islet isolation

Briefly, pancreatic islets were isolated by injection of collagenase (Viacyte CIzyme RI cat. 005-1030) into the pancreatic duct followed by a stationary digestion for 17min in a 37°C drybath. After density centrifugation, islets were hand-picked and cultured overnight in RPMI 1640 containing 5.5 mM glucose, 10% FBS and P/S at 37 °C with 5% CO_2_.

#### Stem cell-derived β cells (sc-β cells) and islet aggregates

##### iPSC culture

Human induced pluripotent stem cells (iPSCs) were obtained from an individual with T1D, as published previously^27^. Rnls exon 2 was targeted for deletion using a dual gRNA strategy in undifferentiated iPSCs and gene knockdown was verified by qPCR. Prior to starting differentiations, iPSCs were cultured in 2D planar form in 75cm^2^ tissue culture-treated flasks (Celltreat cat. 229341) coated with GelTrex (Fisher cat. A1413302; 1h at 37°C) and maintained in mTeSR media (Stem Cell Technologies cat. 85850). Rock inhibitor 10μM Y-27632 was added at seeding and after passages.

##### Differentiation to sc-β cells

iPSCs were differentiated into sc-β cells using the Hogrebe et al. protocol ^82^. All growth factors and media compositions are the same as the listed protocol. At stage 0 (S0), 0.6-0.8 x10^5^ cells/cm^2^ were seeded in 6 well tissue culture-treated plates coated with GelTrex. Extra plates were included to measure established quality control (QC) markers using flow cytometry and/or immunohistochemistry (IHC) throughout S0-S6. Stage-specific media was changed late morning at the same time +/- 30mins.

##### Quality control (QC)

In S0-S5, sc-β cell QC by IHC was measured after fixation and permeabilization (BD cat. 554714). ICC buffer (0.1% Triton-X, 5% donkey serum (Jackson Labs cat. 100181-234), PBS) was added for 1h at room temperature (RT) to block and permeabilize cells. After PBS wash, primary antibodies for stage-specific markers were diluted 1:300 in ICC (c-peptide, Developmental Studies Hybridoma Bank (DSHB), University of Iowa, cat. no. GN-ID4-S; nkx6.1, DSHB cat. no. F55A12-S) and added overnight at 4°C. Primary antibodies were aspirated, secondary antibodies (Thermo anti-rat 594 cat. A-11007; anti-mouse cat. A-11001) diluted 1:300 in ICC for 1h and wells washed 3x with PBS. Cells were co-stained with Hoescht 33342 (Thermo cat. H3570; 1:10 000) 10mins. prior to imaging. For flow cytometry at S6, sc-β cells were dispersed using TrypLE, centrifuged, and resuspended in BD fixation buffer for 30min at 4°C. After centrifugation, cells were resuspended in BD permeabilization buffer for 30min at 4°C and blocked with ICC for 30min. Cells were incubated in primary antibodies for c-peptide and nkx6.1 diluted 1:50 in staining buffer (0.5% BSA, 2mM EDTA in PBS) for 30mins at RT. Secondary antibodies were directly added (1:50) and resuspended in staining buffer for 30mins. Cells were centrifuged and resuspended in staining buffer before analysis on BD Fortessa II. Each sample was filtered to reduce cell clumping.

##### Sc-β cell aggregates

On S6 day 7, some planar WT and Rnls KO sc-β cells were used to generate islet-like aggregates. Sc-β cells were dissociated using TrypLE Express (Gibco cat. 12604013) for 5mins at 37°C to generate single cell suspensions. Cells were resuspended to 1 x10^6^ cells/mL in ESFM media, 5mL added to each well of a 6 well plate, and plates transferred to an orbital shaker at 37°C, 5% CO_2_. Aggregates formed over a 7 day period. Media was changed every other day until functional experiments (Seahorse and GSIS, see below) were performed on S6 day 14.

### Method details

#### Bulk RNA sequencing

Bulk RNA sequencing data was revisited for secondary analysis^81^. Briefly, WT and Rnls^mut^ NIT-1 cells were used for RNA isolation using Zymo Quick-RNA miniprep plus kit (Zymo Research cat. R1058). Novogene Corporation Inc. prepared RNA libraries and performed sequencing (Illumina NovaSeq 6000 system). Pathway analysis using Kyoto Encyclopedia of Genes and Genomes (KEGG) and Gene Ontology (GO; Molecular Function and Biological Process) were used to identify the most significantly changed and most significantly upregulated pathways. Expression of rate-limiting genes in critical pathways presented in Figure 1 is displayed as fragments per kilobase of transcript (FPKM) as a normalization of gene length and the total number of mapped reads.

#### qPCR

##### NIT-1 β cells

NIT-1 cells were grown in 25mM DMEM. Media was aspirated and cells scraped into TriZol (Fisher cat. 15596018). RNA was extracted using a commercial kit (Zymo Research cat. 2050) and quantified using a NanoDrop. RNA was reverse transcribed to cDNA (Azura Genomics cat. AZ-1995) per manufacturer’s instructions. Each forward-reverse qPCR primer pair (Table 1) was validated for efficiency against a standard curve (data not shown) prior to measuring samples (Azura Genomics cat. AZ-1910) and ppia was used as a housekeeping control^83^.

##### Mouse islets

Islets from mice were collected and maintained in RPMI overnight to remove acute stress from harvesting protocols. Mice that received PG *in vivo* had islets maintained with 5μM PG. The following day, islets were handpicked into TriZol for RNA extraction and processed for gene expression analysis (as above).

Cell line and islet qPCR were performed using ABI7900 RTPCR (Life Technologies) and fold change was calculated using the ΔΔ*CT* method.

#### Metabolomics

##### Untargeted metabolomics

WT and Rnls^mut^ NIT-1 β cells were seeded in 6-well plates at 3.75x10^5^ cells/well and grown to 70% confluence (36h) in DMEM. Cells were quenched in 75mM ammonium carbonate (pH 7.4), scraped and extracted in ice cold 40:40:20 acetonitrile:methanol:water solvent. Separate wells were extracted and counted to normalize for differences in biomass between cell lines (no significant difference). Extracts were kept at - 80°C for short-term storage until analysis.

For high-throughput untargeted metabolomics, cell extracts were diluted 1:20 in methanol, and 5 μl of each sample was analyzed in two technical replicates by flow-injection, non-targeted metabolomics in negative ionization mode on an Agilent 6550 Quadrupole Time-of-flight mass spectrometer (Agilent Technologies) as previously described^84^. The mobile phase contained isopropanol/water (60:40, v/v) 1 mM ammonium fluoride with a flow rate of 150 μl per min. The mobile phase was spiked with hexakis(1H,1H,3H-tetrafluoropropoxy)phosphazene and taurocholic acid *ad* ∼1x10^5^ signal intensity online mass calibration. Mass spectra were recorded in profile mode from m/z 50 to 1000 with a frequency of 1.4 spectra/s for 0.46 min at 4 GHz (HiRes). Ions were putatively annotated by matching their inferred mass with compounds in the HMDB database, allowing a tolerance of 1 mDa, and only deprotonated ions were included in the adduct list. Pathway enrichment analyses were performed using Metaboanalyst software (https://www.metaboanalyst.ca/) based on KEGG metabolic pathways.

##### [^13^C_6_]-glucose targeted metabolomics

WT and Rnls^mut^ NIT-1 cells were seeded in the same experimental setup as untargeted analysis and maintained in DMEM supplemented with heavy labeled 5mM [^13^C_6_]-glucose for 24h (Cambridge Isotope Laboratories cat. 110187-42-3). Two 6-well plates of WT cells were treated with 5μM PG for the duration of the experiment. Plates were carefully extracted in randomized sequence to maintain precise timing.

Steady-state targeted metabolic tracer measurements were conducted on a QExactive HF-X mass spectrometer equipped with a HESI II probe. The mass spectrometer was coupled to a Vanquish binary UPLC system (Thermo Fisher Scientific, San Jose, CA). For chromatographic separation prior to mass analysis, 5 ul of each metabolite extract was injected into an Intrada OA column (150 mm x 2 mm, 3 um particle size, Imtakt USA). Mobile phase A was 100 mM ammonium formate in 90% water and 10% acetonitrile, and mobile phase B was 1% FA in 90% acetonitrile and 10% water. The column oven was held at 45°C and autosampler at 4°C. The chromatographic separation was performed as described in Koley et al^85^.The mass spectrometer was operated in full-scan negative mode, with the spray voltage set to 3 kV, the capillary temperature to 320 °C, and the HESI probe to 300 °C. The sheath gas flow was set to 40 units, the auxiliary gas flow to 8, and the sweep gas flow to 1 unit. Data acquisition for all samples was performed in full scan mode in range m/z = 70–800, with the resolution set to 120,0000. For a pooled study sample, a PRM experiment was conducted to ensure metabolite identification; inclusion lists were constructed based on observed RT and m/z in the MS1 data and fragmentation voltage was set to N(CE) 30, 50, and 150. Water, acetonitrile, and formic acid were purchased from Fisher and were Optima LC/MS grade. Ammonium formate was purchased from Honeywell Fluka. Metabolites were identified with authentic standards as well as labelling and fragmentation patterns. Raw data were processed using emzed and custom Python scripts^86^. Total signal of each mass isotopolog of interest was corrected for natural abundance of ^13^C and normalized to baseline.

#### Glucose-stimulated insulin secretion (GSIS)

Static GSIS in planar sc-β cells was performed on day 14 of stage 6 exactly according to Hogrebe et al^82^. Planar sc-β cells were seeded in a 24-well plate and treated with 5μM PG overnight. For the assay, media was replaced with Krebs Ringer Buffer (KrB) and supernatants collected in response to 1h 2mM glucose followed by 1h 20mM glucose. Insulin was measured using a commercially available human insulin ELISA (ALPCO cat. 80-INSHU-E01.1) normalized to cell count collected at the end of the experiment.

#### Seahorse

Specific sensor cartridges (Agilent) for all experiments were hydrated >18h at 37°C (no CO_2_).

##### NIT-1 and MIN6

Metabolic function was assayed using Seahorse XFe96 well microplates for cell lines (Agilent cat. 103792-100). Cells were passaged normally and seeded at 1.0x 10^5^ cells/well (MIN6) or 6.0x 10^4^ cells/well (NIT-1) in 200uL normal DMEM media + supplements and maintained at 37°C, 5% CO_2_. Plates were left in the tissue culture hood for 1h, to avoid cell edge effects, and then maintained at 37°C, 5% CO_2_ until the experiment. On experiment day, growth media was carefully aspirated, washed 3x with seahorse XF DMEM media (Agilent cat. 103575-100; without HCO_3_, serum or phenol red), left at 37°C (no CO_2_) for 1h. Media was aspirated and replaced once more before running. Assay-specific metabolic compounds were diluted day-of in XF DMEM media immediately prior to experiments: Oligomycin 5μM (Sigma Aldrich cat. 495455), Rotenone 2.5μM (Sigma Aldrich cat. 557368) or Antimycin A 2.5μM (Sigma Aldrich cat. A8674. In some experiments, NIT-1 and MIN6 were seeded 48h before running Seahorse assays and treated with 5-100μM PG throughout the pre-experiment period; volumes <5uL PG were added to minimize osmotic changes. H_2_O was added as a vehicle and osmotic control. DMEM growth media was carefully aspirated and replaced every 24h + PG.

Both NIT-1 and MIN6 β cell lines are insulinomas. In n=1 experiment, both cell lines were cultured in low glucose DMEM (5mM glucose + L-glutamine + P/S) for 14 days in an attempt to shift cells towards lower basal glycolysis. For both NIT-1 and MIN6, 8.0x10^4^ cells/well were seeded the night prior on a 96-well plate and treated with 5mM 2-deoxy-D-glucose 2h prior to the initial seahorse reading (2-DG; Millipore Sigma cat. D2134). ATP production rates from glycolysis (glycoATP; pmol/min.) and oxidative phosphorylation (mitoATP; pmol/min.) were estimated using Agilent software.

##### Mouse islets

Islet function was measured using Seahorse XFe24 islet capture microplates (Agilent cat. 101122-100). Islets were collected the day prior and maintained in RPMI 1640 media with 11mM glucose, FBS and P/S at 37°C, 5% CO_2_. On the day of Seahorse, islets were inspected for quality, handpicked and washed in Seahorse MA media (Agilent cat. 103575-100) supplemented with 1% heat-inactivated FBS and 3mM glucose. After centrifugation, islets were resuspended to 50 islets per 500μL, distributed evenly across wells, and carefully repositioned into the depressed islet chamber using a 20μL pipette. Islet screens were slowly set in place by forceps. XFe24 well plates were maintained at 37°C (no CO_2_) for 1h to equilibrate to 3mM glucose prior to the assay. Assay-specific metabolic compounds were diluted day-of in MA media immediately prior to experiments (see above).

##### Sc-β cells and aggregates

XFe96 well plates were coated with 40µL GelTrex at 37°C for 1h and aspirated immediately before use. Sc-β cells were detached using TrypLE, counted, and resuspended in 200µL ESFM media for 1.5x 10^5^ cells/well. Plates rested for 1h in the tissue culture hood and were then maintained at 37°C, 5% CO_2_ overnight. Prior to experiments, media was aspirated, and wells washed 3x with MA media. Plates were maintained at 37°C (no CO_2_) for 1h prior to the assay. Assay-specific metabolic compounds were diluted day-of in MA media immediately prior to experiments (FCCP 1μM; Sigma Aldrich cat. C2920).

All seahorse data was analyzed using Seahorse Analytics Software (Agilent).

#### *In vitro* stress experiments

##### NIT-1 and MIN6 seahorse

Plated cells were treated with dithiothreitol (DTT; Millipore sigma cat. 10197777001) – to mimic protein misfolding stress – 1h prior to the baseline seahorse measurement; 5mM DTT for NIT-1 and 1mM for MIN6 based on pilot data on viability and activating transcription 4 (ATF4) abundance (not shown). H_2_O was added as a vehicle and osmotic control. Viability was measured using Trypan Blue.

##### NIT-1 ^13^C_6_ targeted metabolomics

Plated cells were treated with 5mM DTT 1h prior to metabolite extraction. H_2_O was added as a vehicle and osmotic control.

##### Sc-β cell stress outcomes

Round-bottom 96-well plates were coated with GelTrex and cells seeded at a density of 5.0x10^4^ cells/well in ESFM (low glucose) media on S6 day 15. To expand on previous findings showing ER stress protection in Rnls^mut^ cells, we challenged planar sc-β cells with high glucose (HG; 25mM), DTT (10μM) or hydrogen peroxide (H_2_O_2_; 100μM) to mimic hyperglycemia, ER stress and oxidative stress, respectively. PG and/or stressors were added for 48h and plates maintained at 37°C, 5% CO_2_. On experiment day, cells were gently dispersed using TrypLE, manually transferred to a v-shaped 96-well plate for staining, washed 1x with staining buffer, and stained as follows before flow cytometry quantification (BD Fortessa II):

● ThiolTracker: a proxy for intracellular glutathione (GSH). Cells were stained with 20mM ThiolTracker (Thermo cat. T10095) for 30mins at 37°C, washed, and resuspended in staining buffer (0.5% BSA, 2mM EDTA in PBS).
● 2’,7’-dichlorodihydrofluorescein diacetate (DCFDA): a marker of intracellular reactive oxygen species (ROS); 5μM DCFDA (Thermo cat. 6827) for 30mins at 37°C. Cells were washed and resuspended in staining buffer.
● MitoSox: an indicator of mitochondrial superoxide production; 1μM (Thermo cat. M36005) for 30mins at 37°C. Cells were washed and resuspended in staining buffer.
● Annexin V/7-AAD: apoptosis and viability markers (BD Biosciences cat. 556421); stained according to manufacturer’s instructions using specific binding buffer (1:20 Annexin V, 1:100 7-AAD).

#### Western blotting

##### Rnls localization

NIT-1 cells were grown to confluence in a 10cm dish with DMEM media. Cells. were collected and processed for cytosolic, nuclear and mitochondrial fractions using a commercial kit (Abcam cat. 109719; 5mL of starting buffer and 1mL Buffer A + 1mL buffer/C throughout). Lamelli buffer (BioRad cat. 1610747) was added to samples. Proteins were separated on a 10% gel and transferred to nitrocellulose membranes. Membranes were blocked in 5% milk for >1 h at RT and incubated in primary antibodies overnight 4°C. Renalase antibodies Abcam cat. Ab31291 (1:100), NOVUS cat. 98475 (1:500) and Proteintech cat. 60128 (1:500) were directly compared using appropriate HRP-conjugated secondary antibodies.

*NIT-1 ATF4:* NIT-1 cells were NIT-1 cells were grown to confluence in a 10cm dish with DMEM media, collected, and redistributed in a 12-well plate in DMEM. The next day, cells were treated with 0, 5, 10 or 20mM DTT for 1h. Cells were detached manually by pipette mixing. A small 10μL sample was removed for measuring cell viability (Trypan Blue). Remaining cells were centrifuged, resuspended in RIPA buffer (Thermo cat. 89900) supplemented with a protease inhibitor cocktail (Thermo cat. 78440), lysed using a sonicator and incubated at -80°C to burst cell membranes. Protein content was measured using DC Assay and samples processed as above to probe for activating transcription 4 (ATF4) abundance (Cell Signaling cat. 11815) as a marker of ER stress.

##### Mouse islets

After PG treatment *in vivo*, islets were collected as usual and maintained overnight in RPMI. Islets from Akita mice treated with oral PG were maintained in 5μM PG overnight. The following day, islets were handpicked and lysed in RIPA buffer supplemented with protease inhibitor. ATF4 abundance was quantified using the same methods as NIT-1 ATF4 (see above).

All images were obtained using a ChemiDoc (BioRad, V3.0) and Image Lab densitometry software.

#### *In vivo* experiments

##### Monoamine oxidase assay

Pargyline (PG) inhibits both Rnls and the monoamine oxidase (MAO) enzyme family. There is no commercially available enzyme activity assay for Rnls. To determine which oral dose of PG elicits *in vivo* effects, Akita mice were treated with 0-25µg/mL PG for 7 days. Livers, due to high tissue expression of MAO (Human Protein Atlas) and ease of collection, were homogenized in RIPA lysis buffer supplemented with protease inhibitors. Protein content was measured using a DC protein assay (BioRad cat. 5000111) and lysates were equalized with water to load 50µg of protein. MAO activity was measured in luminescence units (RLU) using extra protocol steps specific to MAO-B (Promega cat. V1401); PG has been shown to inhibit this isoform specifically.

##### Oral Pargyline (PG) administration

At day 19-20, Akita littermates were matched for weaning blood glucose and allocated to normal drinking water or water supplemented with 25-50µg/mL PG (Sigma cat. P8013) for 7-28 days, depending on the experiment. Littermate WT mice acted as controls in all studies. Water was changed every other day. We repeated experiments in n=3-6 cohorts of mice to ensure consistency in phenotypes.

##### Blood glucose and insulin

Mice were fasted for 12h overnight. Blood glucose was measured using a glucometer (Contour Next) the following morning. Blood was collected using capillary tubes (Sarstedt cat. 16440100) centrifuged at 10000 x g for 5mins to isolate serum. Serum insulin was measured by commercial ELISA (EMD Millipore cat. EZRMI-13K).

##### Glucose tolerance tests (IPGTT)

To assess whole-body glucose tolerance, mice were administered an intraperitoneal bolus of glucose (2 g/kg) after a 5h fast. Blood glucose was sampled from the tail vein and recorded using a glucometer from 0-120min post-injection. GTTs were performed after 4weeks of PG treatment, and 3-4 days after the 12h overnight fast for insulin sampling. Area under the curve (AUC) was calculated using GraphPad Prism 8 software (GraphPad Software) by subtracting baseline glucose in mg/dL to reflect the true response.

##### Insulin tolerance tests

To assess *in vivo* insulin sensitivity, mice were administered a bolus of insulin (0.75IU/kg; Eli Lilly, Humulin R) after a 2h fast; this duration is meant to standardize blood glucose as best possible without forcing reliance on tissue glycogen stores. Blood glucose was sampled from the tail vein and recorded using a glucometer from 0-90min post-injection. AUC was calculated using GraphPad Prism 8 software (GraphPad Software) by subtracting baseline glucose in mg/dL to reflect the true response.

#### Histology

##### β cell mass

Pancreas were dissected from anesthetized mice, weighed, spread in cassette and fixed in 4% paraformaldehyde (PFA) for 24h for paraffin embedding. Adjacent full footprint sections were immunostained for glucagon (in-house antibody; Dr. Michael Appel) and insulin (Bio-Rad cat. 5330-0104G) using a standard ABC peroxidase protocol (Vector Labs cat. SP-2001, PK-4001, PK-4007, SK-4105). Primary antibodies were incubated overnight at 4°C (insulin 1:500, glucagon 1:3000). Akita mice are characterized by altered islet mantle architecture and degranulation of insulin so both stains were necessary to distinguish β cell versus α cell mass. β cell mass was calculated by point counting morphometry using methods previously established^87^.

## Data Analysis & Statistics

Data are presented as mean ± SEM. Each dot represents an individual mouse or pooled n=3-5 technical replicates from independent experiments in cell culture-derived data. Sc-β cell data are the average of n=2-3 independent differentiations. Flow cytometry data were analyzed with FlowJo software v.10.6.1 (FlowJo LLC) and y-axes were normalized to mode. Statistical significance was determined in GraphPad Prism 10 software. One or two-way ANOVA with Holm Sidak posthoc testing for multiple comparisons were used in most experiments; in datasets with comparisons between two groups (i.e. viability and untargeted metabolomics data), student’s t-test with Welch’s correction was used.

## Acknowledgements

We thank Jen Hollister-Lock and Brooke Sullivan for their expertise, teaching and gracious guidance on islet dissections and histology, and Christopher Cahill for help with tissue preparation.

This work was supported by NIH RO1 (grant #, S.K and P.Y), Mary K. Iacocca Foundation (T.M), Joslin Diabetes Research Center (2P30DK036836), the Joslin Islet Isolation Core, and the Diabetes Research and Wellness Foundation Chair (S.B.W).

## Author contributions

T.L.M, S.K, G.C.W, S.B.W and P.Y designed animal studies. T.L.M and B.M conducted all genotyping. T.L.M and B.R designed, conducted and analyzed all metabolomics experiments. S.K, P.Y, G.C.W and S.B.W supervised the project. E.C and Y.I generated NIT-1 cell lines. S.W helped conduct all iPSC to sc-β cell differentiations. J.A.D.S.P analyzed sc-β cell death/apoptosis and helped with c-peptide/NKX6.1 staining. T.L.M wrote the manuscript with input from all authors. G.C.W and S.B.W contributed significant and meaningful discussion throughout.

---

**Extended Data 1:**
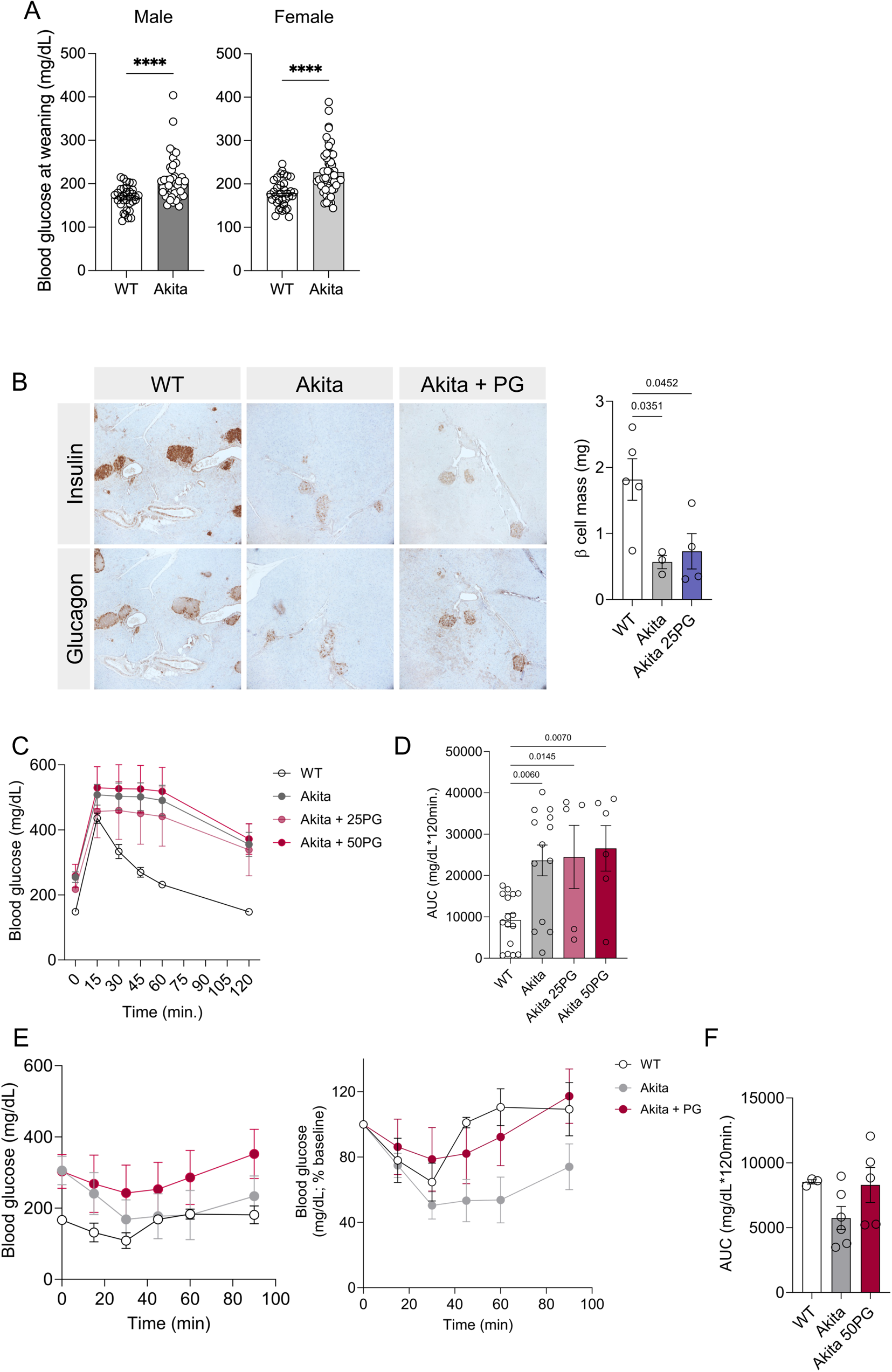
(A) Blood glucose at weaning for male and female WT and Akita mice in all experiments; n=33-54/group. (B) Insulin and glucagon staining in male pancreas; n=3-5/group. (C) GTT in female mice at 4 weeks. The GTT area under the curve (AUC) is shown in (D); n= 6-13/group. (E) ITT in female mice, also expressed as % change from baseline in (F; no significant comparisons). Data is expressed as mean ± SEM. Statistical significance indicated by full p-value comparison or with *** p ≤ 0.0001 as indicated. Each dot represents an individual mouse.

**Extended Data 2:**
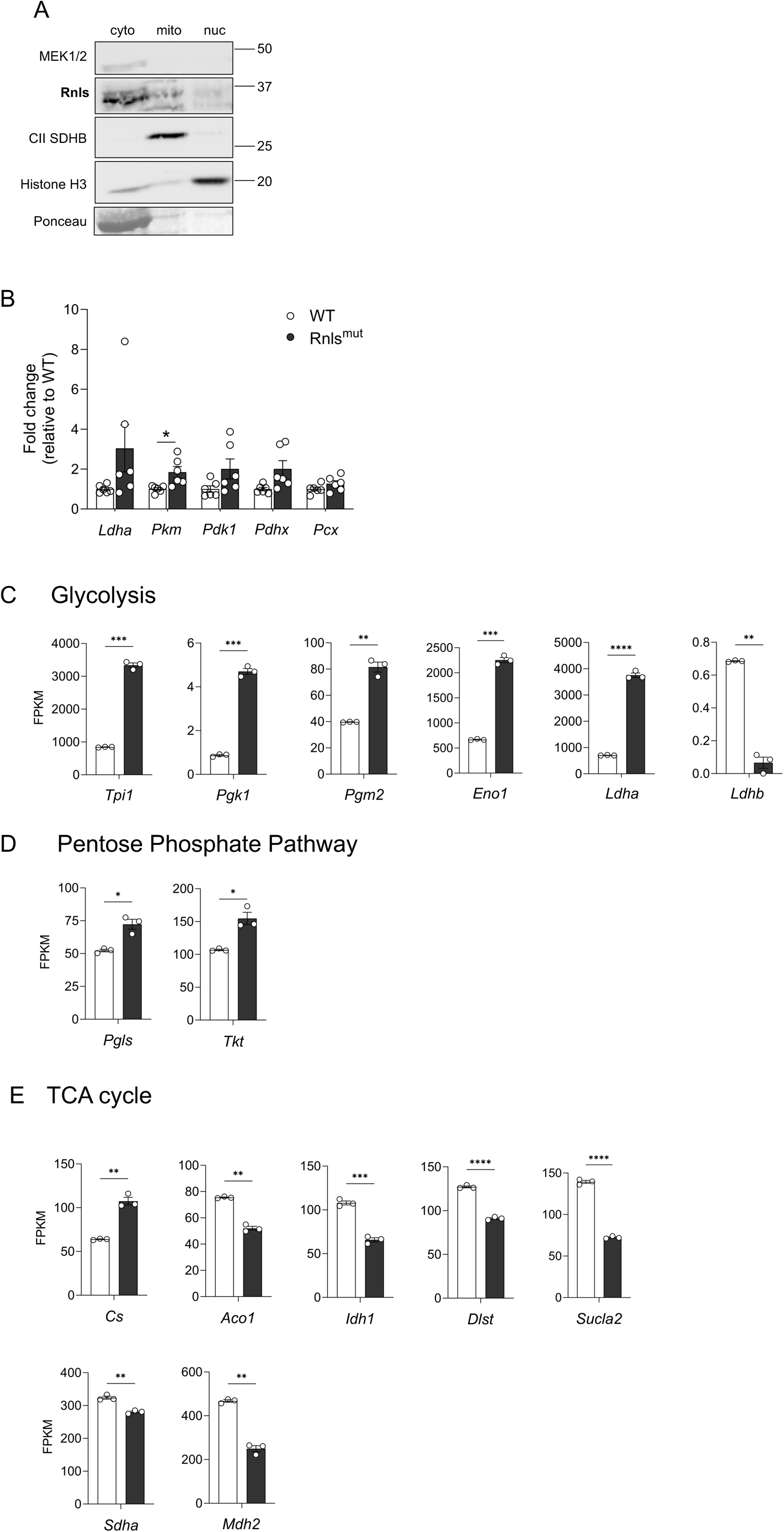
Rnls is located in the cytoplasm of β cells and regulates core metabolic pathways. (A) Western blotting for subcellular localization of Rnls protein in NIT-1 β cells: MEK1/2 is cytosolic, CII (SDHB) is mitochondrial, Histone H3 is nuclear and ponceau red staining is shown for total protein. (B) qPCR confirmation of key glucose metabolism transcripts identified in RNAseq. Expression of enzymes related to (C) Glycolysis; (D) Pentose phosphate pathway or (E) TCA cycle from RNAseq. Data is expressed as mean ± SEM; n= 3-6 technical replicates. Statistical significance indicated by *p ≤ 0.05; ** p ≤ 0.01; *** p ≤ 0.001; *** p ≤ 0.0001 as indicated. Abbreviations: tpi1, triosephosphate isomerase 1; pgk1, phosphoglycerate kinase 1; pgm2, phosphoglucomutase 2; eno1, enolase 1; ldha, lactate dehydroganase a; ldhb, lactate dehydrogenase b; pgls/6PGL, 6-phosphogluconolactonase; g6pdx, glucose-6-phosphate dehydrogenase; tkt, transketolase; cs, citrate synthase; aco1, aconitase 1; idh1, isocitrate dehydrogenase 1; dlst, 2-oxoglutarate dehydrogenase complex, mitochondrial; sucla2, succinate-CoA ligase ADP-forming subunit beta; suclg1, succinate-CoA ligase GDP/ADP-forming subunit alpha; sdha, succinate dehydrogenase complex flavoprotein subunit A; mdh2, malate dehydrogenase 2.

**Extended Data 3:**
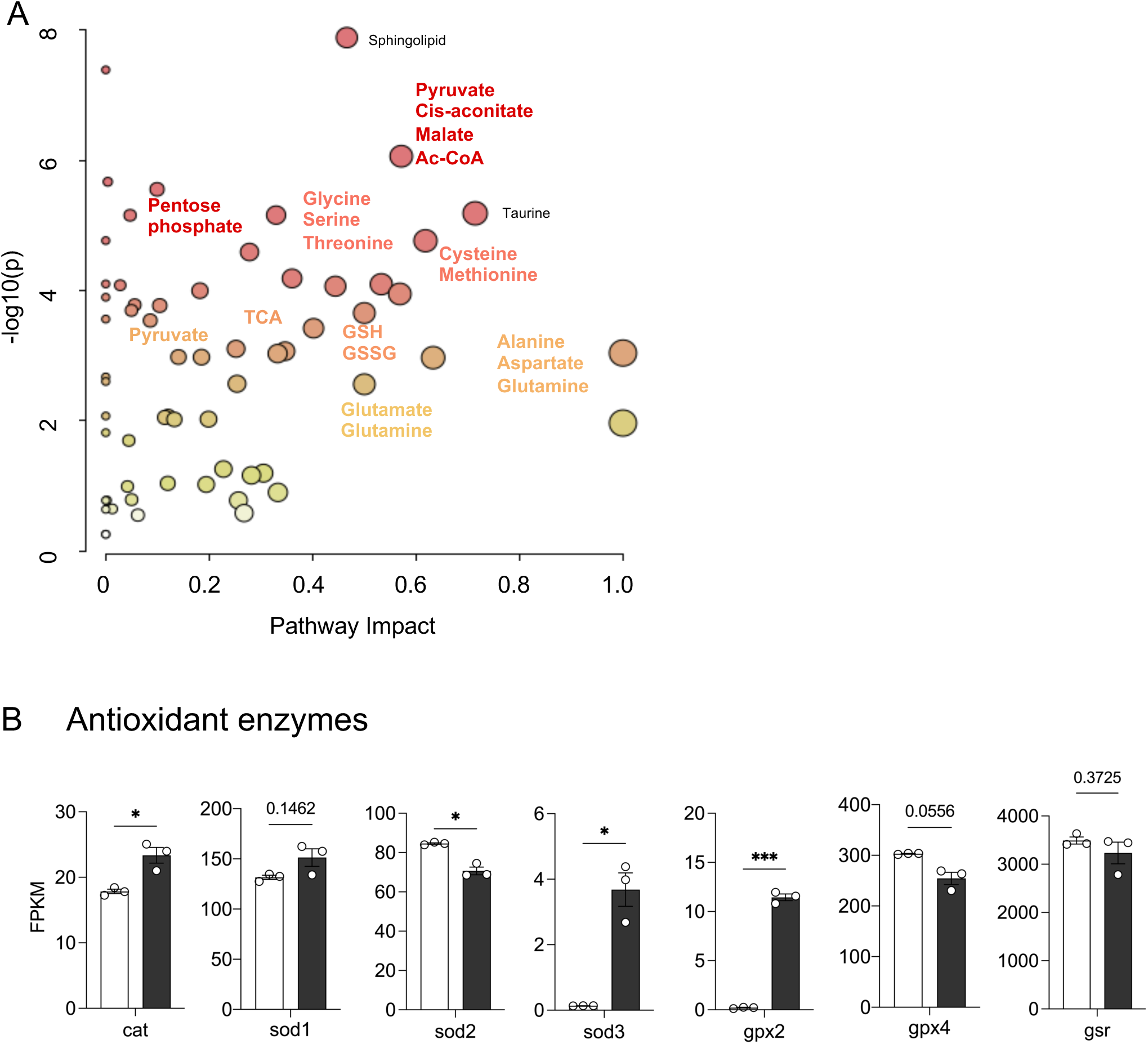
(A) KEGG pathway enrichment in Rnls^mut^ NIT-1 cells determined by Metaboanalyst software. (B) Expression of key antioxidant enzymes from RNAseq analysis. Data is expressed as mean ± SEM. Statistical significance indicated by *p ≤ 0.05; *** p ≤ 0.001; as indicated. Abbreviations: cat, catalase; sod1, superoxide dismutase 1; sod2, superoxide dismutase 2; sod 3, superoxide dismutase 3; gpx2, glutathione peroxidase 2, gpx4, glutathione peroxidase 4; gsr, glutathione disulfide reductase.

**Extended Data 4:**
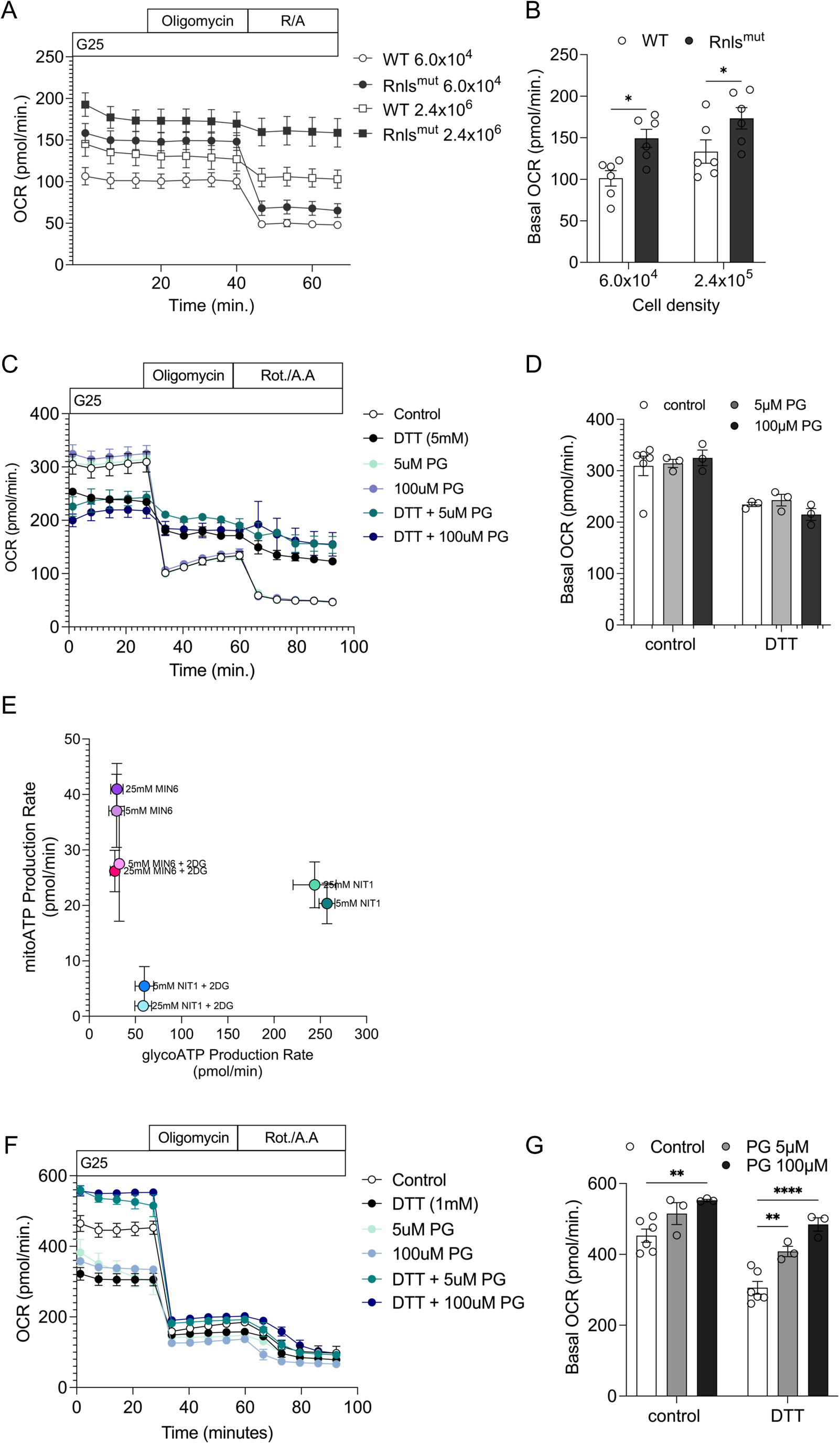
Mitochondrial metabolism is less affected by Rnls inhibition at rest and during stress. Oxygen consumption rate (OCR) in (A) WT versus Rnls^mut^ NIT-1 β cells, with basal OCR quantified in (B). Data is n=6 technical replicates from one representative experiment; repeated n=3 times in WT vs Rnls^mut^ NIT-1 β cells. NIT-1 cells treated with Pargyline (PG) for 24h and challenged with DTT for 1h in (C). Basal OCR is quantified in (D). (E) Shows ATP produced from glycolytic (glycoATP) versus mitochondrial (mitoATP) energy metabolism pathways in cells cultured in 5mM or 25mM glucose DMEM media, in the absence or presence of 5mM 2-DG to test glycolytic inhibition in NIT-1 vs MIN6 cells. MIN6 cells treated with 24h PG and challenged with DTT for 1h in (F) and basal ECAR quantified in (G). Data is n=3-6 technical replicates from one representative experiment; repeated n=2 times in WT vs PG-treated MIN6. Data is expressed as mean ± SEM. Statistical significance indicated by *p ≤ 0.05; ** p ≤ 0.01; *** p ≤ 0.001; *** p ≤ 0.0001 as indicated.

**Extended Data 5:**
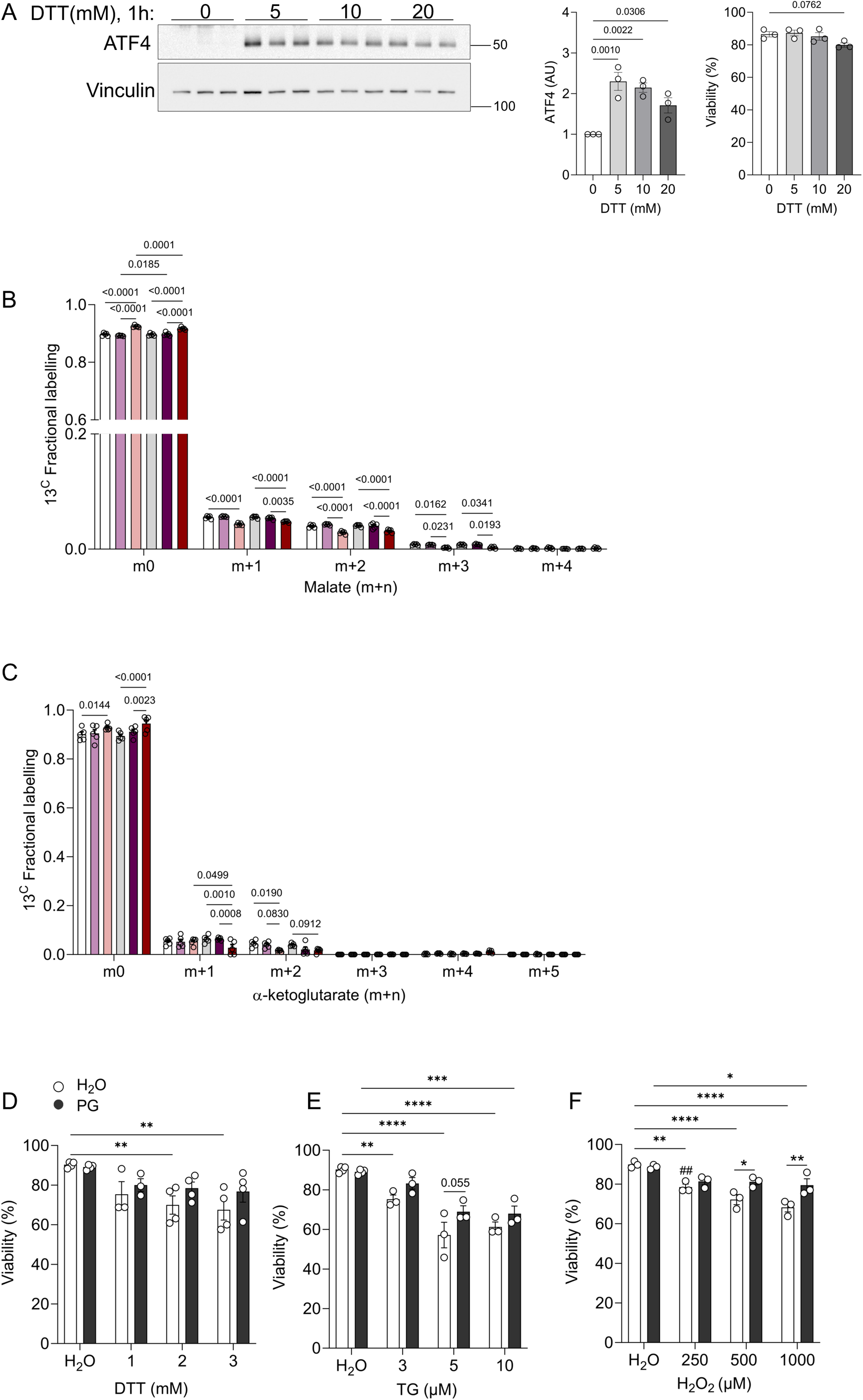
(A) Wildtype NIT-1 cells were treated with 0-20mM DTT for 1h to measure ER stress pathway activation via ATF4 protein abundance. Vinculin is included as a total protein loading control. Cell viability was measured in an aliquot of the same cells using Trypan blue staining. (B) Panels and (C) show labelling of [^13^C_6_]-glucose into malate and α-ketoglutarate m+0 to m+n isotopologs. Viability (% total) of NIT-1 β cells pre-treated with 5µM PG and exposed to (D) DTT, (E) TG or (F) H_2_O_2_ stress for 48h. Data is expressed as mean ± SEM; each dot represents a biological replicate, n=3-5. Statistical significance indicated by full p-value comparisons or *p ≤ 0.05; ** p ≤ 0.01; *** p ≤ 0.001; *** p ≤ 0.0001 as indicated.

**Extended Data 6:**
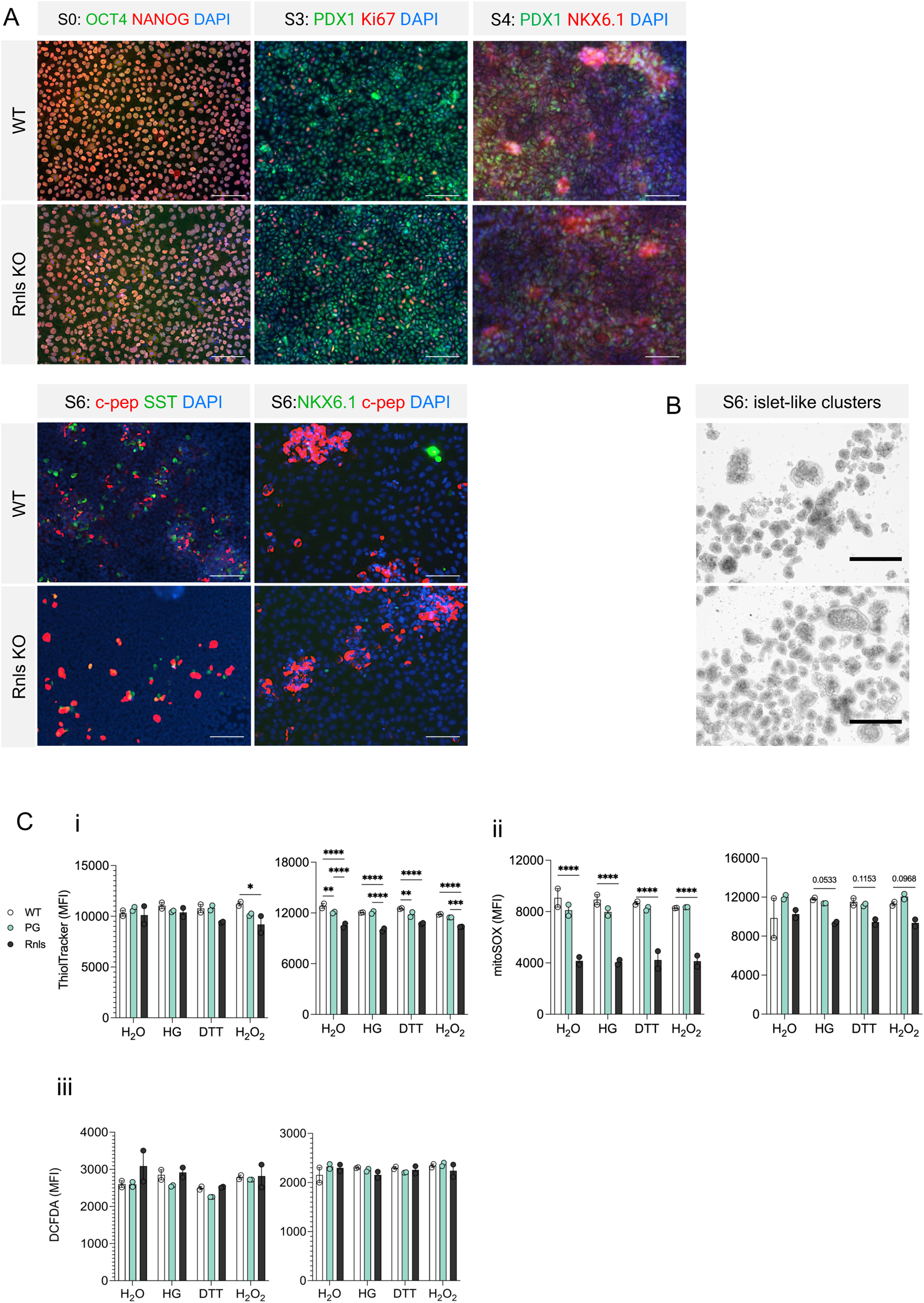
(A) Immunohistochemistry for quality control of WT and Rnls KO iPSC lines (S0, stage 0) through differentiation steps to mature (S6) mature sc-β cells with stage-specific markers. (B) Aggregated islet-like clusters formed at the end of S6. (C) Quantifications for Fig 5G; n=2 experiments measuring (i) ThiolTracker (GSH); (ii) mitoSOX (mitochondrial ROS) and (iii) DCFDA (cytosolic ROS) in sc-β cells in response to stress. Data are expressed as mean ± SEM. Statistical significance indicated by full p-value comparison as indicated.

**Source Data (supports Figure 1):**
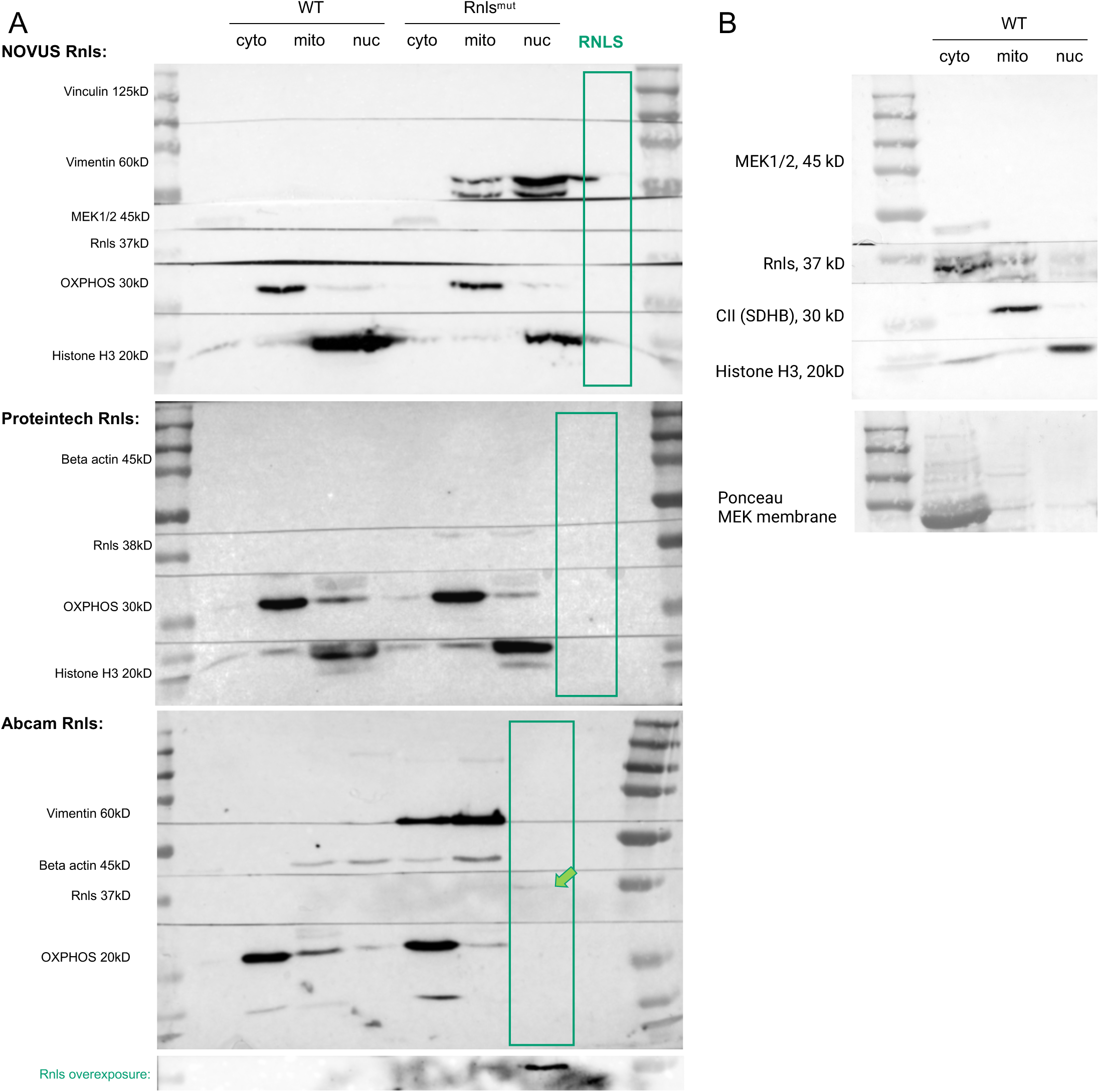
(A) Western blotting for Rnls protein in NIT-1 β cells. Three different commercial antibodies were used: NOVUS, Proteintech and Abcam with purified RNLS as a control. In (B), subcellular cytosolic (cyto), mitochondrial (mito) and nuclear (nuc) fractions were probed. Full gels are shown; n=1 experiment.

## Notes

### Competing Interest Statement

The authors have declared no competing interest.

